# *Gl*Rac regulates an ATG8-independent autophagy-like pathway in *Giardia lamblia*

**DOI:** 10.64898/2026.02.17.706457

**Authors:** Angélica Hollunder Klippel, Corryn Newman-Boulle, Grant Reed, Dule Dule, Saanya Kedia, Martin T Kelty, Wai Pang Chan, Han-Wei Shih, Alexander R Paredez

## Abstract

Autophagy is a conserved catabolic process essential for cellular homeostasis and adaptation to nutrient stress. The protozoan parasite *Giardia lamblia* lacks most canonical autophagy-related (ATG) genes, including the hallmark ATG8, raising longstanding questions about whether this deeply divergent parasite can perform autophagy. Here, we identify an ATG8-independent autophagy-like pathway in *Giardia* regulated by *Gl*Rac, the parasite’s sole Rho family GTPase. *Gl*Rac-positive double-membrane compartments are induced by encystation and nutrient depletion, and their abundance rapidly declines following amino acid replenishment but is unaffected by glucose, indicating amino acid-specific regulation. *Giardia* Target of Rapamycin (GTOR) levels decrease during nutrient depletion, and GTOR knockdown increases compartment abundance, identifying GTOR as a negative regulator of compartment formation and linking this pathway to nutrient sensing. Time-lapse microscopy revealed that these compartments form through linear and cup-shaped intermediates before becoming spherical and are subsequently cleared upon nutrient replenishment. Of nine putative ATG orthologs examined, none localized as specifically as *Gl*Rac to these structures, supporting the existence of a highly divergent pathway. Nevertheless, the compartments exhibit multiple conserved autophagy-associated features, including double-membrane morphology, actin recruitment, acidification, and cysteine protease activity. Pharmacological inhibition of cysteine proteases with E-64d or of V-ATPase-mediated acidification with concanamycin A promotes compartment accumulation, consistent with continuous degradative turnover. *Gl*Rac regulates compartment biogenesis bidirectionally: constitutive activation increases compartment abundance and size, whereas knockdown reduces them. Finally, quinacrine, an FDA-approved antigiardial drug that accumulates in acidic organelles, perturbs *Gl*Rac-positive compartments, consistent with its reported effects on autophagy in other eukaryotes, raising the possibility that this pathway contributes to parasite fitness. Together, these findings establish *Gl*Rac as a central regulator of an ATG8-independent autophagy-like pathway in *Giardia* and demonstrate that this parasite retains key structural, regulatory, and degradation-associated features of autophagy despite the apparent absence of most canonical ATG machinery.

**Author Summary:** Autophagy is a conserved cellular process that helps cells adapt to nutrient limitation by remodeling and degrading their own components. The intestinal parasite *Giardia lamblia*, which causes widespread diarrheal disease, lacks recognizable homologs of most genes associated with canonical autophagy, raising the question of whether this highly divergent parasite can perform the process at all. We found that *Giardia* forms previously uncharacterized membrane-bound compartments in response to nutrient depletion and during differentiation into its infectious cyst stage. These compartments share key features associated with autophagosomes, including double membranes, acidification, and enzymes involved in protein degradation. We also identified a regulatory protein called *Gl*Rac as a central regulator of these compartments. Similar proteins regulate autophagy in plants, fungi, and animals, suggesting that this regulatory mechanism is broadly conserved across eukaryotes. Finally, quinacrine, a drug historically used to treat giardiasis, promotes the accumulation of these compartments, consistent with its reported effects on autophagy in other eukaryotes. Together, our findings reveal that *Giardia* retains a highly divergent autophagy-like pathway despite the apparent absence of much of the recognizable canonical autophagy machinery, providing new insight into the evolution of autophagy and the cell biology of this important human pathogen.

## Introduction

Macroautophagy, hereafter referred to as autophagy, is a conserved catabolic process by which eukaryotic cells degrade and recycle cytoplasmic components. This pathway involves the formation of a double-membrane phagophore cup that engulfs cytoplasmic cargo and then matures into a sealed autophagosome. The double-membrane autophagosome fuses with lysosomes or vacuoles to enable the enzymatic degradation and recycling of its contents. By removing damaged components and recycling nutrients, autophagy maintains cellular homeostasis and supports adaptation to various stressors. Given its pivotal role in responding to nutrient limitation, a challenge likely faced by both ancient and modern eukaryotes, autophagy is thought to have emerged in the last eukaryotic common ancestor (1–4). In protozoan parasites, complex life cycles and fluctuating environments have shaped the retention and functional specialization of autophagy (5,6).

Across eukaryotes, over 40 ATG proteins have been identified (7,8), with about 20 forming the conserved core machinery required for autophagosome biogenesis (9). In contrast to *Saccharomyces cerevisiae* and mammals, many parasitic protists possess a restricted subset of ATGs that still support autophagosome formation and degradation despite lacking several canonical components (5,10,11). For instance, components of the ATG1 kinase complex are highly divergent or absent in some species, and key mediators of membrane expansion, such as ATG2 and ATG9, are missing in others. Notably, several protists also lack the ATG12 conjugation system required for phagophore expansion (5,6,10,12). These observations suggest the existence of alternative or streamlined autophagic pathways in these organisms. One of the most striking reductions in ATG protein content is observed in *Giardia lamblia* (syn. *G. intestinalis*, *G. duodenalis*), a microaerophilic protozoan parasite and early-branching eukaryote characterized by a highly compact genome and reduced cellular complexity (13–15). This parasite causes giardiasis, a major cause of waterborne diarrhea worldwide that affects both humans and other mammals, with an estimated 280 million human infections annually (14,16,17). *Giardia* has a biphasic life cycle, consisting of two distinct forms: a dormant, environmentally resistant cyst and a replicative trophozoite. Infection begins when cysts are ingested, followed by excystation in the small intestine, where trophozoites emerge and colonize the intestinal epithelium (13). To complete the cycle and enable transmission, trophozoites undergo encystation, a differentiation process that generates infective cysts capable of surviving outside the host (18).

Comparative genomics has revealed a limited set of core autophagy genes in *Giardia*, including ATG1, VPS15, VPS34, ATG16, and ATG18, as well as the upstream regulator TOR (5,10,11,19). Notably, the parasite lacks genes encoding the ATG8 and ATG12 conjugation systems, which are essential for phagophore expansion and autophagosome maturation in canonical autophagy pathways (5,6,10,11). This reduced set of ATGs in *Giardia* aligns with the overall simplification of its cellular machinery, which includes a streamlined metabolic repertoire and the absence of canonical organelles such as mitochondria, Golgi apparatus, endosomes, and lysosomes (15,20,21). In particular, the lack of classical lysosomes raises questions about how autophagic degradation could occur in this parasite (5).

However, despite the absence of classical lysosomes, *Giardia* possesses a peripheral vacuole (PV) system comprising numerous small vacuoles located underneath the plasma membrane. These compartments function in endocytosis and intracellular degradation and have been proposed to perform lysosome-like functions in the parasite (20,22,23). Early cytochemical studies identified acid phosphatase activity within PVs, supporting their classification as lysosome-like organelles (24). Consistent with this role, PVs are acidic, as demonstrated by the accumulation of acidotropic probes including acridine orange and LysoTracker, and contain additional hydrolytic enzymes (25,26). Among these, certain cathepsin-like cysteine proteases have been localized to these vacuoles (26–28). Notably, despite its reduced cellular organization, the *Giardia* genome encodes 27 cathepsin-like cysteine proteases (29). Although the functions of many remain unknown, characterized members of this family have been implicated in parasite differentiation, including encystation and excystation, as well as in virulence through disruption of the host epithelial barrier, degradation of immune mediators, and modulation of intestinal inflammatory responses (27–33).

Together, the reduced repertoire of canonical ATG proteins, the absence of classical lysosomes, and the highly streamlined cellular organization of *Giardia* contributed to the longstanding assumption that the parasite lacks a functional autophagy pathway (5,6,10,11,19). Nevertheless, previous studies have suggested the presence of autophagy-like processes in *Giardia*. Treatment with sterol biosynthesis inhibitors such as azasterols was reported to induce autophagosome-like structures in trophozoites (34). Similarly, exposure to metronidazole analogs induced morphological and ultrastructural changes characteristic of autophagy, including multilamellar concentric membranes (35). Additional evidence came from a study on programmed cell death, in which trophozoites displayed autophagy-like responses to starvation stress (19). In that context, cells accumulated vacuoles labeled with monodansylcadaverine (MDC), a nonspecific marker of autophagic activity (36). However, the presence of double-membrane compartments was not confirmed, the quality of MDC staining was poor, and concerns about cell integrity after prolonged incubation in phosphate-buffered saline have raised questions about the reliability of these results. More recently, a large-scale protein localization study identified numerous GFP-positive structures classified as “putative inclusions,” which were not associated with known organelles. While their identity remains uncertain, the authors noted that some of these structures may represent previously undescribed endomembrane compartments rather than protein aggregation artifacts (37). Overall, the lack of rigorous quantification, appropriate controls, and validated markers has hindered the acceptance of these studies as definitive evidence of autophagy in *Giardia*.

Despite increasing evidence for autophagy-like processes in *Giardia*, little is known about the molecular mechanisms that regulate this pathway. The parasite encodes a single Rho family GTPase, *Gl*Rac, which has been implicated in regulating the actin cytoskeleton and membrane-associated cellular processes (38,39). Notably, Rho GTPases have emerged as important regulators of autophagy across diverse eukaryotes. In *Arabidopsis*, the Rho family GTPase ROP8 promotes autophagosome formation, whereas in mammalian cells, RhoA positively regulates starvation-induced autophagy (40,41). Given its unique status as the sole Rho family GTPase in *Giardia* and its established involvement in cytoskeletal and membrane remodeling, *Gl*Rac is a compelling candidate regulator of autophagosome biogenesis in this parasite.

Using cellular imaging, genetic perturbation, and pharmacological tools, we characterized the formation and properties of these autophagosome-like organelles and identified *Gl*Rac as a key regulator of their biogenesis. Our findings provide compelling evidence for an ATG8-independent autophagy-like pathway in *Giardia*, highlighting both the evolutionary adaptability of this essential process and its potential as a therapeutic target in parasitic protists.

## Results

### *Gl*Rac marks a double-membrane organelle induced by encystation

During our investigation into the role of *Giardia*’s sole Rho family GTPase, *Gl*Rac (GL50803_008496), in the encystation process, we observed its localization to structures with variable morphology, including linear, cup-shaped, and spherical forms (Fig 1A). To determine whether these structures are induced by encystation, we quantified their frequency over a 4 h time course using a strain with integrated Halo-*Gl*Rac (S1 Fig). The proportion of cells containing Halo-*Gl*Rac-labeled structures (cup-shaped and spherical) increased from 13% at 0 h to approximately 30% after 3 h of encystation induction (Fig 1B), indicating that these structures form in response to the differentiation stimulus. Among positive cells, the number of structures per cell was similar at 0 h and 4 h, indicating that encystation increases the proportion of cells forming these compartments rather than their number per cell. In both conditions, most cells contained one Halo-*Gl*Rac-labeled structure, with fewer containing two.

**Fig 1.**
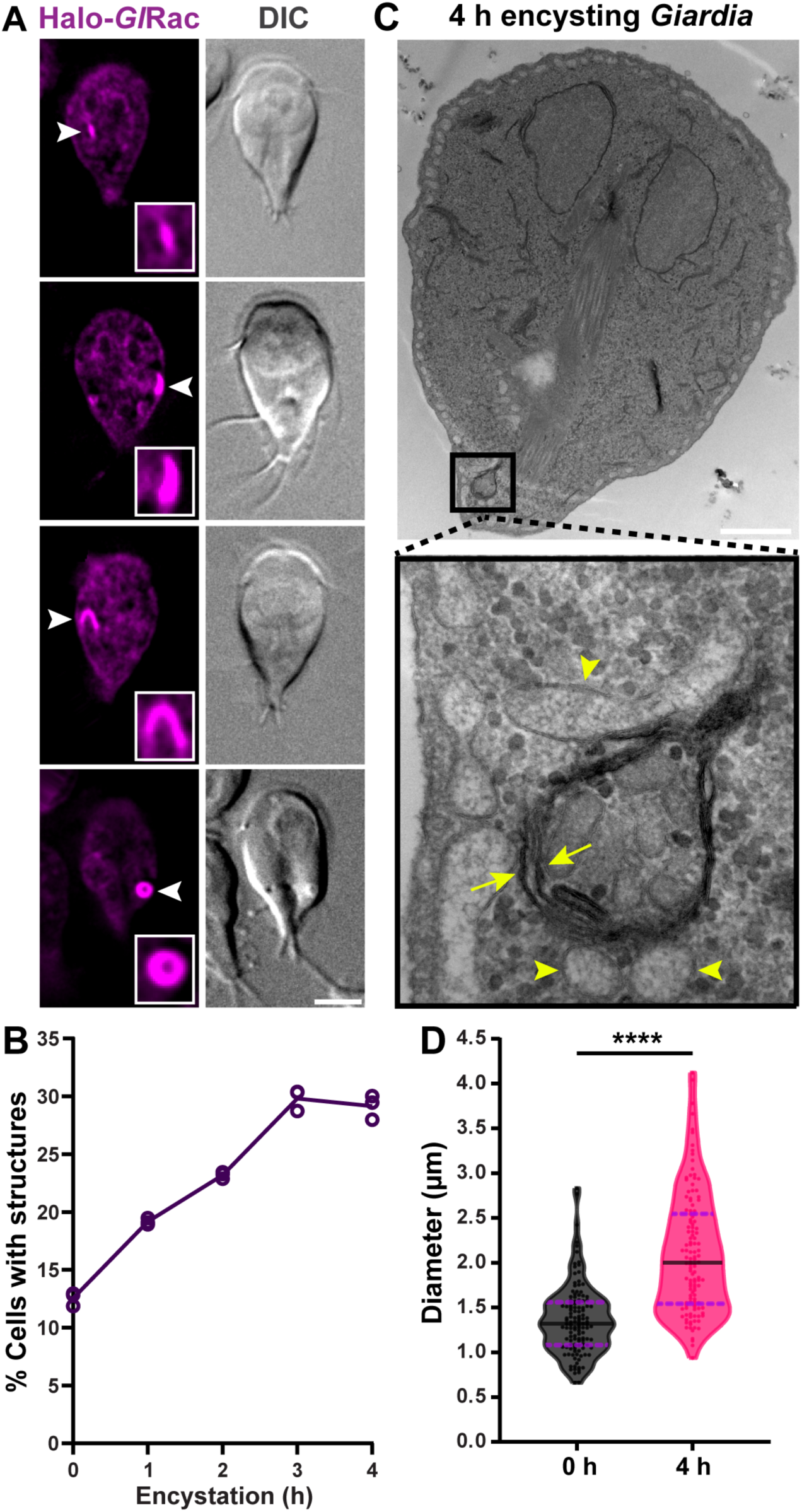
*Gl*Rac localizes to a double-membrane organelle induced by encystation. **(A)** Localization of Halo-*Gl*Rac 4 h post-induction of encystation. Halo-*Gl*Rac concentrates in cellular structures with variable morphology (white arrowheads). Scale bar, 5 µm. **(B)** Quantification of cells harboring Halo-*Gl*Rac-labeled structures (cup-shaped or spherical) at 0, 1, 2, 3, and 4 h post-induction of encystation. For each time point, n > 1,000 cells. Data represent three independent experiments. **(C)** TEM of a 4 h encysting cell showing a double-membrane (yellow arrows) compartment, consistent with an autophagosome-like organelle. Yellow arrowheads indicate potential peripheral vacuoles surrounding the compartment. Scale bar, 1 µm. **(D)** Violin plot showing the size distribution (maximum diameter) of spherical *Gl*Rac-labeled compartments measured in compartment-positive cells at 0 h and 4 h after encystation induction (all compartments from n = 100 cells across three independent experiments). Each dot represents one measurement. The solid black line marks the median, and the dashed purple lines indicate the first and third quartiles. P-value was calculated using two-tailed Welch’s t-tests (****P ≤ 0.0001).

To further characterize these structures, we performed transmission electron microscopy (TEM) on wild-type trophozoites after 4 h of encystation. TEM revealed organelles with a double-membrane configuration (Fig 1C), a morphological hallmark of autophagosomes (42,43).

The size of spherical *Gl*Rac-positive compartments, quantified from live-cell image data, significantly increased during encystation. Maximum diameters expanded from an average of 1.36 µm (range: 0.66-2.83 µm) in non-encysting cells to 2.10 µm (range: 0.94-4.12 µm) after 4 h of encystation induction (Fig 1D). Notably, these values exceed the typical sizes reported for autophagosomes in mammalian cells (~0.5-1.5 µm) and in *S. cerevisiae* (~0.4-0.9 µm) (44,45).

The identification of these organelles in encysting trophozoites is notable; their induction during encystation aligns with the well-established role of autophagy in cellular remodeling during differentiation in model organisms such as *S. cerevisiae*, *Dictyostelium discoideum*, and *Caenorhabditis elegans* (3). Similarly, autophagy supports metabolic and morphological changes during host transitions in other protozoan parasites, including *Leishmania* spp., *Plasmodium* spp., and *Trypanosoma cruzi* (5,6,46–48). These functional roles in different systems support the idea that *Giardia* may also rely on an autophagy-related process during differentiation, although the pathway appears highly divergent and reduced.

### Nutrient availability inversely correlates with the number of cells harboring *Gl*Rac-positive structures

Having identified *Gl*Rac-positive organelles in *Giardia* and linked their formation to encystation, we next investigated whether their abundance responds to changes in nutrient availability. Given the well-documented role of starvation in inducing autophagy across diverse organisms (49–52), we hypothesized that *Giardia* may respond similarly to nutrient limitation.

To test this, we quantified the frequency of *Gl*Rac-positive structures in Halo-*Gl*Rac cells cultured for 48, 72, and 96 h (Fig 2A). The proportion of cells containing at least one *Gl*Rac-positive structure rose sharply with prolonged culture and nutrient depletion, increasing from 12.7% at 48 h to 74.7% at 72 h and reaching 91.3% at 96 h (Fig 2B). Among positive cells, the number of *Gl*Rac-positive structures also increased over time (Fig 2C): at 48 h, nearly all positive cells contained a single structure (range: 1-2). By 72 h, two *Gl*Rac-positive structures per cell became the most common outcome (range: 1-4), and this pattern persisted at 96 h, although with a broader distribution (range: 1-5). These observations indicate that nutrient depletion promotes both the generation and accumulation of *Gl*Rac-positive compartments.

**Fig 2.**
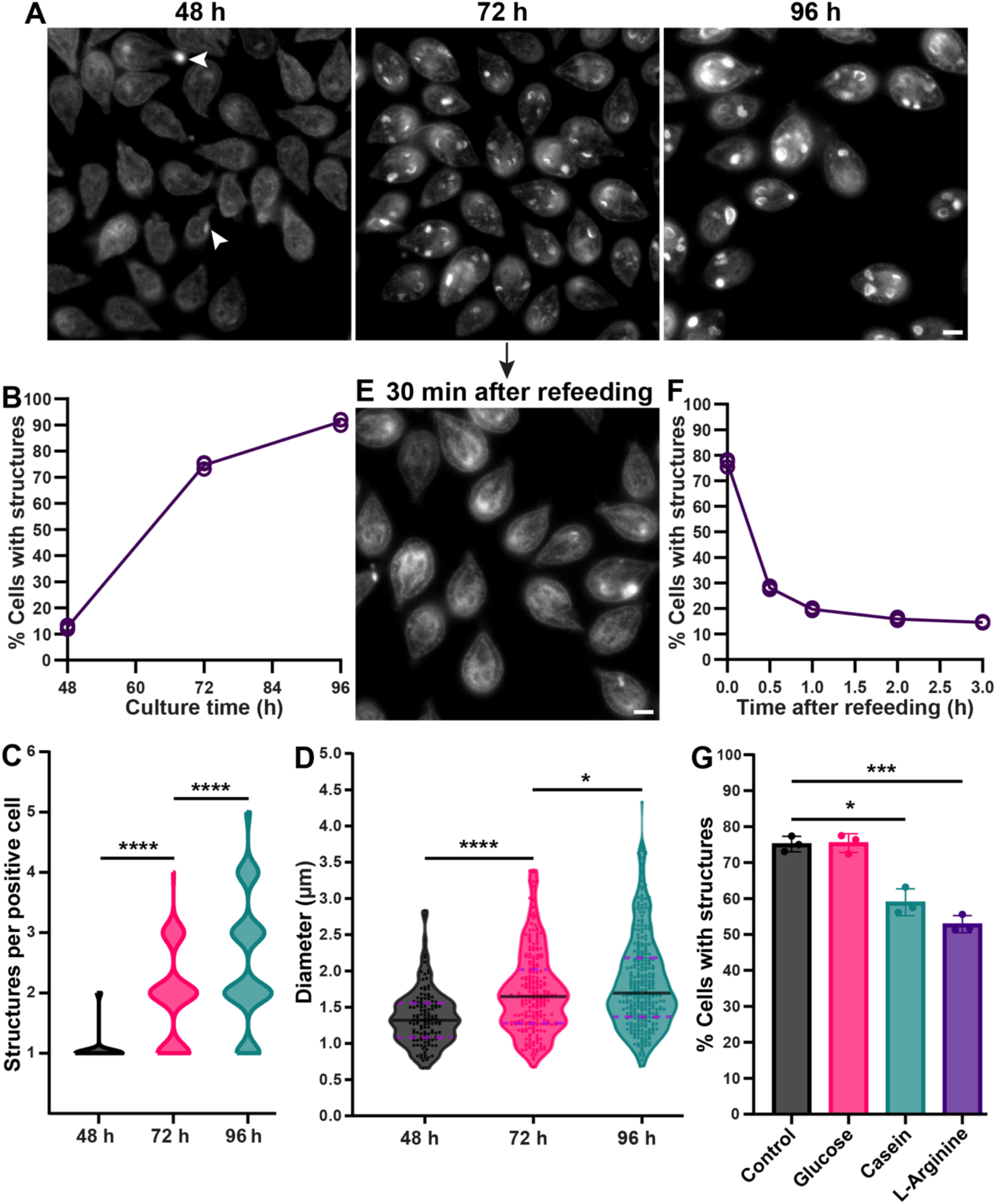
Nutrient availability inversely correlates with the number of cells harboring *Gl*Rac-positive structures. **(A)** Representative images of Halo-*Gl*Rac (gray) cells cultured for 48, 72, and 96 h. White arrowheads indicate Halo-*Gl*Rac-labeled structures at 48 h. **(B)** Quantification of cells containing Halo-*Gl*Rac-labeled structures at the indicated time points. For each time point, n > 1,000 cells. Data represent three independent experiments. **(C)** Violin plot showing the distribution of the number of *Gl*Rac-positive structures in positive cells (cells containing at least one structure) in 48, 72, and 96 h cultures (n = 105 cells from three independent experiments). P-values were calculated using two-tailed Welch’s t-tests (****P ≤ 0.0001). **(D)** Violin plot showing the size distribution (maximum diameter) of spherical Halo-*Gl*Rac-labeled compartments measured in 48, 72, and 96 h cultures (all compartments from n = 100 cells across three independent experiments). Each dot represents one measurement. The solid black line marks the median, and the dashed purple lines indicate the first and third quartiles. P-values were calculated using two-tailed Welch’s t-tests (*P ≤ 0.05; ****P ≤ 0.0001). **(E)** Representative image of Halo-*Gl*Rac cells cultured for 72 h and then refed with fresh medium for 30 min. Scale bar, 5 µm. **(F)** Quantification of cells containing Halo-*Gl*Rac-labeled structures after refeeding for 0.5, 1, 2, or 3 h. For each time point, n > 1,000 cells. Data represent three independent experiments. **(G)** Quantification of cells containing Halo-*Gl*Rac-labeled structures after 72 h of culture and subsequent 2 h supplementation with TYDK salts (Control) or TYDK salts supplemented with 10 g/L glucose, 20 g/L casein digest, or 10 mM L-arginine. For each condition, n > 875 cells. Data are presented as mean ± SD from three independent experiments. P-values were calculated using paired two-tailed t-tests (*P ≤ 0.05; ***P ≤ 0.001).

Nutrient limitation also influenced compartment size (Fig 2D). The diameter of spherical compartments increased from an average of 1.36 µm (range: 0.66-2.83 µm) at 48 h to 1.70 µm (range: 0.68-3.39 µm) at 72 h and 1.84 µm (range: 0.68-4.33 µm) at 96 h. Although on average slightly smaller than those formed during encystation, these compartments fell within a comparable size range.

To determine whether the compartments formed under nutrient limitation share the double-membrane organization characteristic of those generated during encystation, wild-type cells were examined by TEM after 72 h of culture (S2 Fig). These organelles displayed the expected double-membrane organization and also included multilamellar forms (S2B Fig), matching the ultrastructural features observed during encystation (S2A and 1C Figs).

To determine whether starvation-induced structures are reversible, replenishing 72 h cultures with fresh medium reduced the frequency of *Gl*Rac-positive structures to 28.1% after 30 min and returned to near baseline levels after 3 h (Fig 2E and F). This demonstrates that the abundance of *Gl*Rac-positive structures is strongly influenced by nutrient availability, increasing during nutrient depletion and decreasing upon nutrient restoration.

Since amino acid and glucose depletion have been linked to autophagy induction in protozoan parasites (53–56), we sought to determine which nutrients modulate this process in *Giardia*. To test this, we examined whether glucose or amino acids (casein) at concentrations equivalent to standard *Giardia* medium affect *Gl*Rac-positive structures in starved cultures. Although casein digest is not the only amino acid source in the medium, it is the only fully peptide-based component. Arginine was tested separately because it serves as a major energy source and promotes growth at 10 mM (57). Glucose supplementation had no measurable effect, whereas casein digest and L-arginine reduced the frequency of *Gl*Rac-positive structures from 75.2% to 58.9% and 52.8%, respectively (Fig 2G). These results indicate that amino acid availability plays a critical role in regulating *Gl*Rac-positive compartment levels in *Giardia*.

### GTOR suppresses *Gl*Rac-positive compartment formation under nutrient-rich conditions

The response of *Gl*Rac-positive structures to nutrient depletion and amino acid refeeding suggested that their abundance is regulated by nutrient-sensing pathways. In many eukaryotes, the serine/threonine kinase Target of Rapamycin (TOR) plays a central role by sensing nutrient levels, particularly amino acids, and inhibiting autophagy when nutrients are abundant (6,58,59). *Giardia* encodes a putative TOR ortholog, GTOR (GL50803_0035180) (19,60), raising the question of whether this pathway similarly regulates *Gl*Rac-labeled compartment formation in the parasite.

To explore this possibility, we endogenously tagged GTOR with a C-terminal mNeonGreen (mNG) tag in the Halo-*Gl*Rac background and examined its localization under nutrient-rich (24 h) and nutrient-depleted (72 h) conditions. GTOR appeared diffusely distributed in the cytoplasm and did not colocalize with *Gl*Rac-labeled structures (Fig 3A). However, GTOR expression was markedly reduced in 72 h cultures compared to 24 h cultures (Fig 3B), suggesting that GTOR abundance decreases during nutrient limitation.

**Fig 3.**
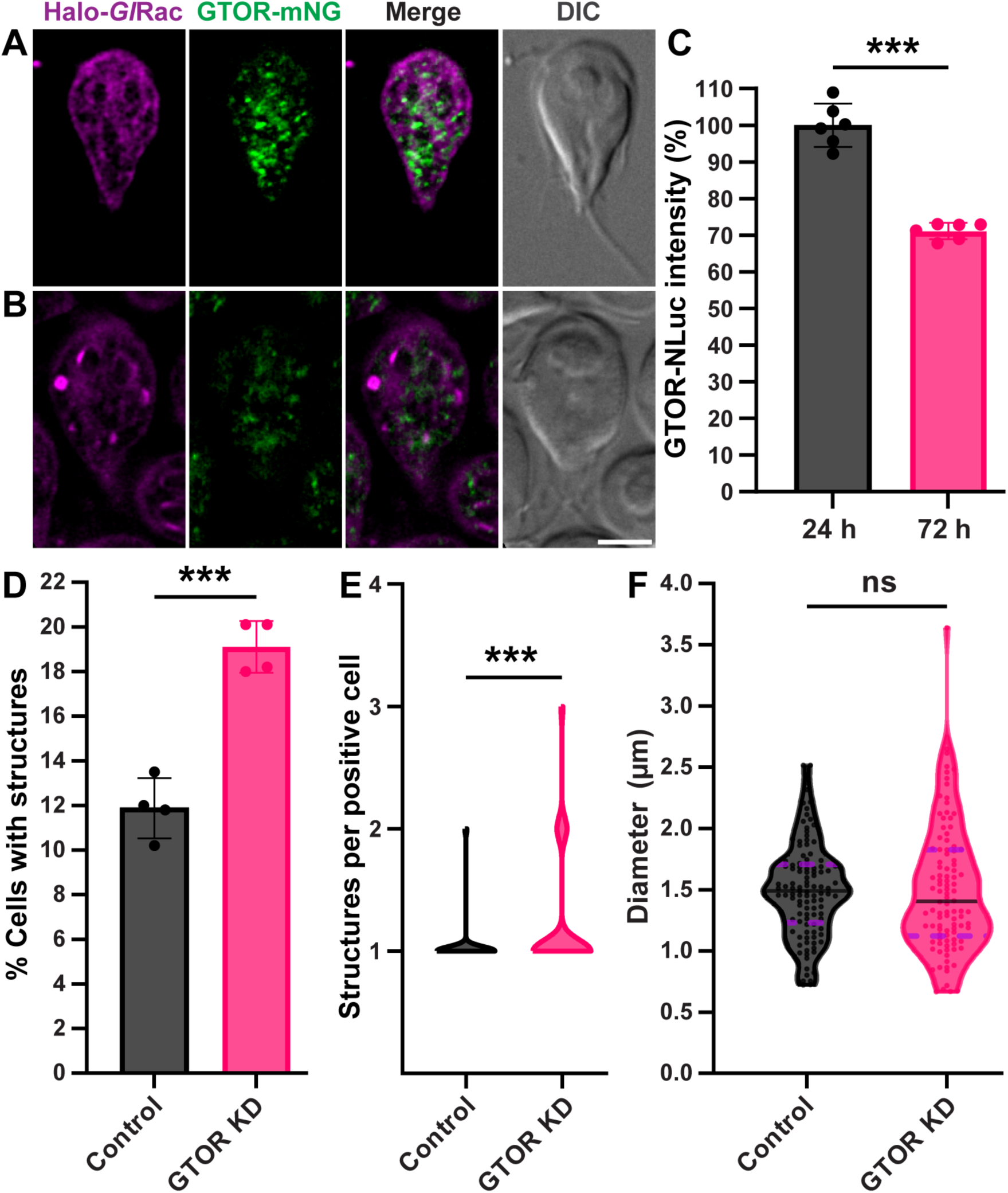
Nutrient depletion reduces GTOR levels, and GTOR negatively regulates *Gl*Rac-positive compartment formation. **(A-B)** Representative images showing the cellular distribution of GTOR (GL50803_0035180-mNG; green) in the Halo-*Gl*Rac (magenta) strain cultured for 24 h **(A)** or 72 h **(B)**, illustrating reduced GTOR abundance in nutrient-depleted cells. GTOR images were acquired using the same exposure settings and processed with identical intensity scaling. Scale bar, 5 µm. **(C)** Relative expression of endogenously tagged GTOR-NLuc in 24 h and 72 h cultures. Data are presented as mean ± SD from six replicates across two independent experiments. P-values were calculated using paired two-tailed t-tests (***P ≤ 0.001). **(D)** Quantification of Halo-*Gl*Rac cells containing *Gl*Rac-labeled structures after 24 h of transfection with either a translation-blocking morpholino targeting GTOR (GTOR KD) or a nonspecific control (control). For each condition, n > 700 cells. Data represent mean ± SD from four independent experiments. P-values were calculated using paired two-tailed t-tests (***P ≤ 0.001). **(E)** Violin plot showing the distribution of the number of *Gl*Rac-labeled structures in positive cells (cells containing at least one structure) in control and GTOR KD conditions (n = 140 cells from four independent experiments). P-values were calculated using two-tailed Welch’s t-tests (***P ≤ 0.001). **(F)** Violin plot showing the size distribution (maximum diameter) of spherical compartments measured in control and GTOR KD conditions (all compartments from n = 100 cells across four independent experiments). Each dot represents one measurement. The solid black line marks the median, and the dashed purple lines indicate the first and third quartiles. P-values were calculated using two-tailed Welch’s t-tests (ns, not significant, P > 0.05).

Because TOR inhibition commonly induces autophagy in other eukaryotes, we next tested whether rapamycin, a potent TOR inhibitor and well-characterized autophagy inducer (58,61), stimulates the formation of *Gl*Rac-labeled structures in *Giardia*. Treatment of 48 h cultures with 36 µM rapamycin for 2 h, a concentration previously shown to affect encystation in *Giardia* (62), did not significantly alter the proportion of cells containing these structures (15.2% vs. 13.8% in controls; S3 Fig). This result suggests that TOR signaling in *Giardia* may be insensitive to rapamycin or that the drug does not effectively inhibit GTOR under the conditions tested.

To test the role of GTOR directly, we depleted GTOR with a translation-blocking morpholino and quantified *Gl*Rac-positive structures under log-growth conditions. GTOR knockdown was confirmed by reduced GTOR-NLuc signal, corresponding to a 69.9% decrease relative to control cells (S4 Fig). GTOR depletion significantly increased the proportion of cells containing *Gl*Rac-labeled structures by approximately 60% relative to control (Fig 3D). GTOR knockdown also increased the number of structures per positive cell, expanding the observed range from 1–2 structures in control to 1–3 structures following GTOR depletion (Fig 3E). In contrast, GTOR depletion did not significantly alter the size of *Gl*Rac-positive structures (Fig 3F), with mean diameters of 1.50 µm in both control and GTOR-depleted cells.

Together with the reduction in GTOR levels observed in nutrient-depleted cultures, these findings indicate that GTOR suppresses the abundance of *Gl*Rac-positive compartments and provide direct functional evidence that TOR signaling regulates this process in *Giardia*.

### Dynamics of *Gl*Rac-positive organelle formation and clearance

Building on the evidence that *Gl*Rac labels compartments with hallmarks of autophagosomes, we next explored how these organelles form and are cleared. Autophagy is a dynamic process involving sequential steps: membrane nucleation, phagophore expansion, membrane curvature, and closure to form a double-membrane autophagosome. This organelle matures and fuses with lysosomes (or with the vacuole in yeast and plants), where acid hydrolases degrade the cargo for recycling (1,9,63).

To investigate the dynamics of *Gl*Rac-positive compartment formation in *Giardia*, we performed time-lapse microscopy using Halo-*Gl*Rac-expressing cells. We observed a sequence of morphological transitions similar to those described in model organisms during autophagosome formation (Fig 4A and B). These events occurred in the ventral region near the bare area (Fig 4A, S1 Movie) and in the dorsal region of the cell (Fig 4B, S2 Movie). The bare area, a microtubule-free region at the center of the ventral disc, has been implicated in membrane trafficking (64). In ventral events, a persistent *Gl*Rac-enriched ventral region was frequently observed adjacent to the site of compartment formation (Fig 4A, S1 Movie). To further characterize this region, we examined the localization of the *Giardia* KDEL receptor (*Gl*KDELR; GL50803_004502-mNG), a conserved component of the ER protein retention machinery (65). *Gl*KDELR was enriched in the same ventral region and colocalized with 81.5% of *Gl*Rac-labeled compartments (S5 Fig). Based on the enrichment of this ER-associated protein, we refer to this *Gl*KDELR-enriched region beneath the ventral disc as the ventral ER.

**Fig 4.**
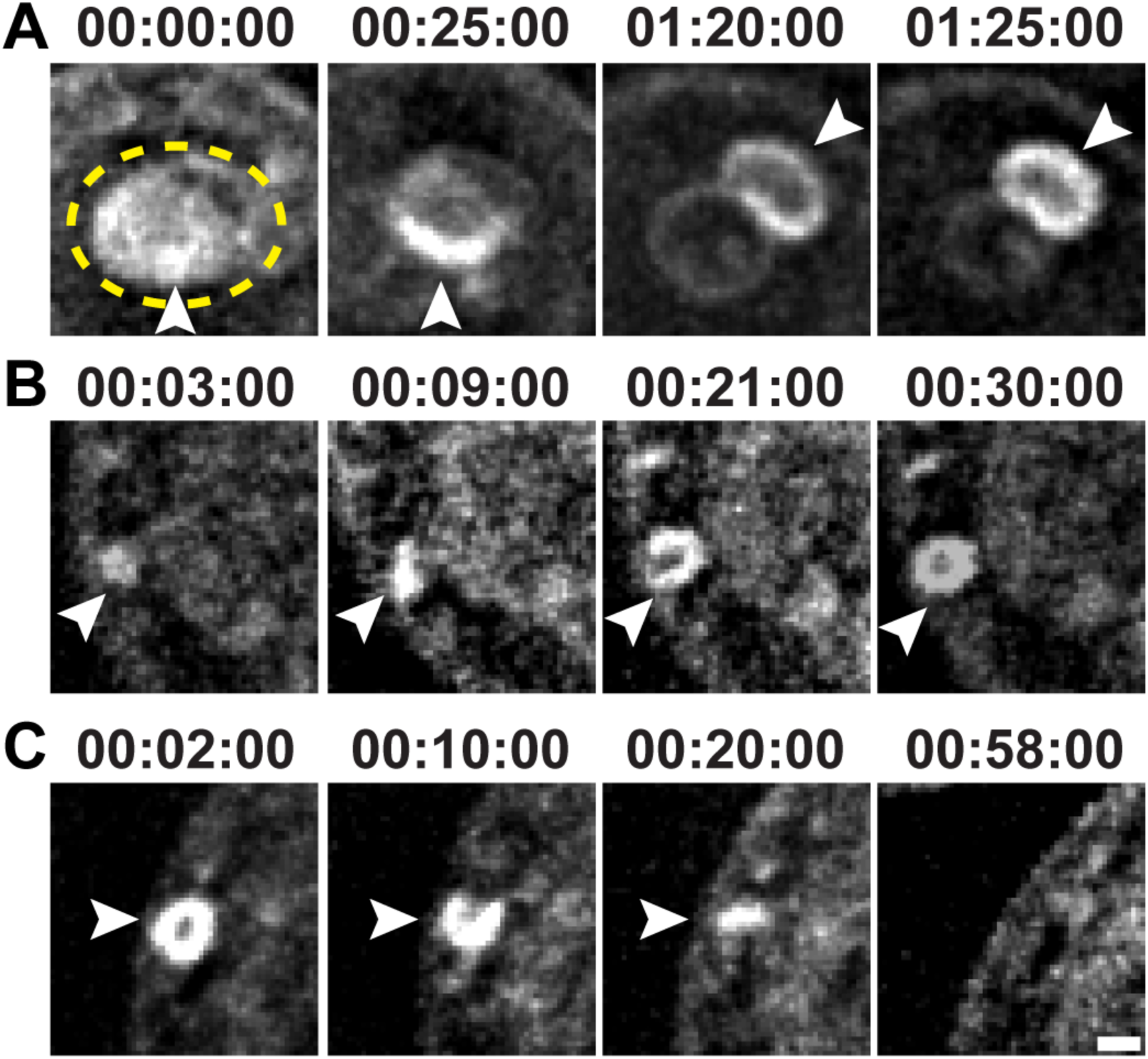
Time-lapse microscopy reveals the dynamics of *Gl*Rac-positive organelle biogenesis and clearance. **(A-B)** Halo-*Gl*Rac (gray)-expressing cells were cultured for 48 h, exposed to nutrient-depleted medium prepared from 96 h cultures, and imaged for 12 h. The yellow dashed circle indicates the ventral region of *Gl*Rac enrichment. Images for B were taken from a compartment forming in the dorsal area. **(C)** Halo-*Gl*Rac-expressing cells were cultured for 72 h, exposed to fresh medium, and then imaged for 12 h. White arrowheads indicate *Gl*Rac-labeled structures. Images represent at least 10 similar events per condition across three independent experiments. Scale bar, 1 µm.

In both regions, *Gl*Rac-positive organelle biogenesis initiated with a discrete focus of Halo-*Gl*Rac accumulation, consistent with a membrane nucleation event. This was followed by an elongation phase during which the compartment maintained a linear morphology, indicative of phagophore expansion. Subsequently, the membrane curved to form a cup-shaped structure resembling the engulfment stage described in other systems, and ultimately developed into a spherical structure, likely representing a fully closed autophagosome (Fig 4A and B) (9,63). In the ventral region, full closure occurred over a range of 1 h 15 min to 6 h 50 min, whereas in the dorsal area, completion was observed as early as 30 min but could take up to 8 h. In contrast, autophagosome biogenesis in *S. cerevisiae* and mammalian cells is considerably faster, typically occurring within 10 min under starvation conditions (66–68). The slower formation of *Gl*Rac-positive compartments in *Giardia* may reflect its reduced repertoire of conserved ATG proteins.

During compartment clearance, the sequence of morphological transitions was reversed (compare panels B and C in Fig 4), taking between 50 min and 4 h 50 min. Spherical compartments reverted to a cup shape, then to linear structures that gradually shrank and were reabsorbed by the cell (Fig 4C, S3 Movie). This reversibility suggests that *Giardia* maintains a regulated mechanism for *Gl*Rac-positive organelle turnover.

In contrast to lysosome reformation in mammalian cells, which involves the emergence of tubules from autolysosomes to regenerate the lysosomal pool (69), we did not observe the formation of tubular protrusions during *Gl*Rac-positive compartment clearance. However, the cup and linear-shaped structures may be tubular (Fig 4C, S3 Movie). Further investigation into the molecular regulators and kinetics of these transitions will provide insights into the unique features of this pathway.

### Cellular localization of putative ATG proteins

Given the similarity between the biogenesis of *Gl*Rac-positive organelles and autophagosomes in model organisms, we next investigated the cellular localization of *Giardia’s* putative ATG proteins. Previous bioinformatic analysis of the *G. lamblia* genome identified a limited set of ATG candidates, including ATG1, ATG18, ATG16, VPS15, and VPS34 (5,19). However, components like VPS15 and VPS34 also participate in other membrane-trafficking pathways and may perform non-autophagic roles in this parasite (5,11). Beyond these in silico predictions, none of these candidates has been experimentally examined in *Giardia*, and their subcellular localization and potential contribution to autophagy remain undetermined.

To begin addressing this, we C-terminally tagged the full set of candidates proposed in earlier surveys and expressed them in the Halo-*Gl*Rac background (S6 Fig). Because these candidates were drawn from the published literature, tagging preceded our own sequence analysis; we subsequently performed reciprocal BLAST searches for each and report the complete analysis, together with the naming conventions used here, in S1 Dataset. Although ATG16 is a conserved component of the phagophore expansion machinery in many organisms, it is also known to be dispensable for autophagy in some protists (6,70). Notably, the top reciprocal BLAST hit for the *Giardia* ATG16 candidate is human POC1, a centriolar protein. Together with its axonemal localization in *Giardia*, this suggests that the candidate is unlikely to function as an ATG-like protein in this organism. Frame-by-frame analysis revealed reproducible signal coincidence between many *Gl*ATG candidates and *Gl*Rac-positive compartments; however, because each candidate was broadly distributed rather than enriched at these sites, we did not interpret this coincidence as robust colocalization (S6 Fig). This inconclusive spatial association underscores the need for functional validation, such as knockdown or overexpression experiments, to determine whether these proteins participate in the autophagic-like response in *Giardia*.

Because a specific *Giardia* protein, GL50803_0028994, has been proposed to function as an ATG8-like protein (62), we tested this claim directly. This protein was reported to colocalize and interact with Myeloid Leukemia Factor (Mlf) vesicles, which are upregulated during encystation and have been suggested to represent autophagosome-like structures involved in protein clearance (62). To assess whether GL50803_0028994 localizes to the *Gl*Rac-positive organelles identified in this study, we tagged it with mNG and expressed it in the Halo-*Gl*Rac strain. GL50803_0028994-mNG overlapped with Halo-*Gl*Rac-labeled structures in only 20.3% of cells and was primarily observed as diffuse cytoplasmic puncta, with no clear enrichment at *Gl*Rac-positive compartments (S7A Fig).

Structural analysis further challenges the classification of GL50803_0028994 as an ATG8 ortholog. Compared to *S. cerevisiae* ATG8 (*Sc*ATG8), the *Giardia* protein lacks the four β-strands that form the ubiquitin-like core required for ATG8 function (71,72) (S7B Fig), yielding a high root-mean-square deviation (RMSD, a standard measure of structural similarity in which lower values indicate greater structural resemblance) of 11.851 Å relative to *Sc*ATG8. This structural divergence is consistent with the weak E-value (2.7 × 10^−2^) reported for this protein in previous bioinformatic analyses (19). Together, these features indicate that GL50803_0028994 is unlikely to be an ATG8 ortholog. Moreover, a recent study of GL50803_0028994’s interaction partner, Mlf, reported cytoplasmic localization without detectable membrane association based on immunogold labeling. In addition, no putative autophagy factors were identified among its interacting proteins, arguing against Mlf and GL50803_0028994 being involved in autophagy (73).

Finally, we examined an ATG7 ortholog candidate (GL50803_006288; E-value = 3.5 × 10^−2^) identified in the same bioinformatic screen (19). Although ATG7 functions as the E1-like enzyme in both the ATG8 and ATG12 conjugation systems (74), the *Giardia* protein localized exclusively to the nuclei and did not colocalize with Halo-*Gl*Rac-labeled structures (S7C Fig). Notably, earlier studies used more stringent thresholds (E-value < 10^−4^) to identify conserved ATGs (10,11). Although these high E-value candidates are unlikely to represent functional ATG8 and ATG7 orthologs, this does not exclude the possibility that *Giardia* encodes highly divergent ATG proteins that escape detection by sequence-based methods while still retaining conserved functions.

### Conserved autophagy machinery is recruited to *Gl*Rac-positive compartments

Considering the reduced number of putative *Gl*ATGs in *Giardia* and their inconclusive colocalization with *Gl*Rac-positive organelles, we next investigated whether other conserved components of the autophagy machinery are recruited to these structures. Actin dynamics play essential roles in autophagy, from autophagosome biogenesis to their fusion with lysosomes and subsequent clearance (75–78). To investigate whether actin associates with these newly identified organelles, we C-terminally tagged actin (GL50803_0040817) with Halo and expressed it in the mNG-*Gl*Rac strain. Under nutrient-depleted conditions, actin-Halo was robustly recruited to mNG-*Gl*Rac-labeled compartments in 88.9% of cells containing these structures (Fig 5A and I), including during the early stages of phagophore formation (Fig S8).

**Fig 5.**
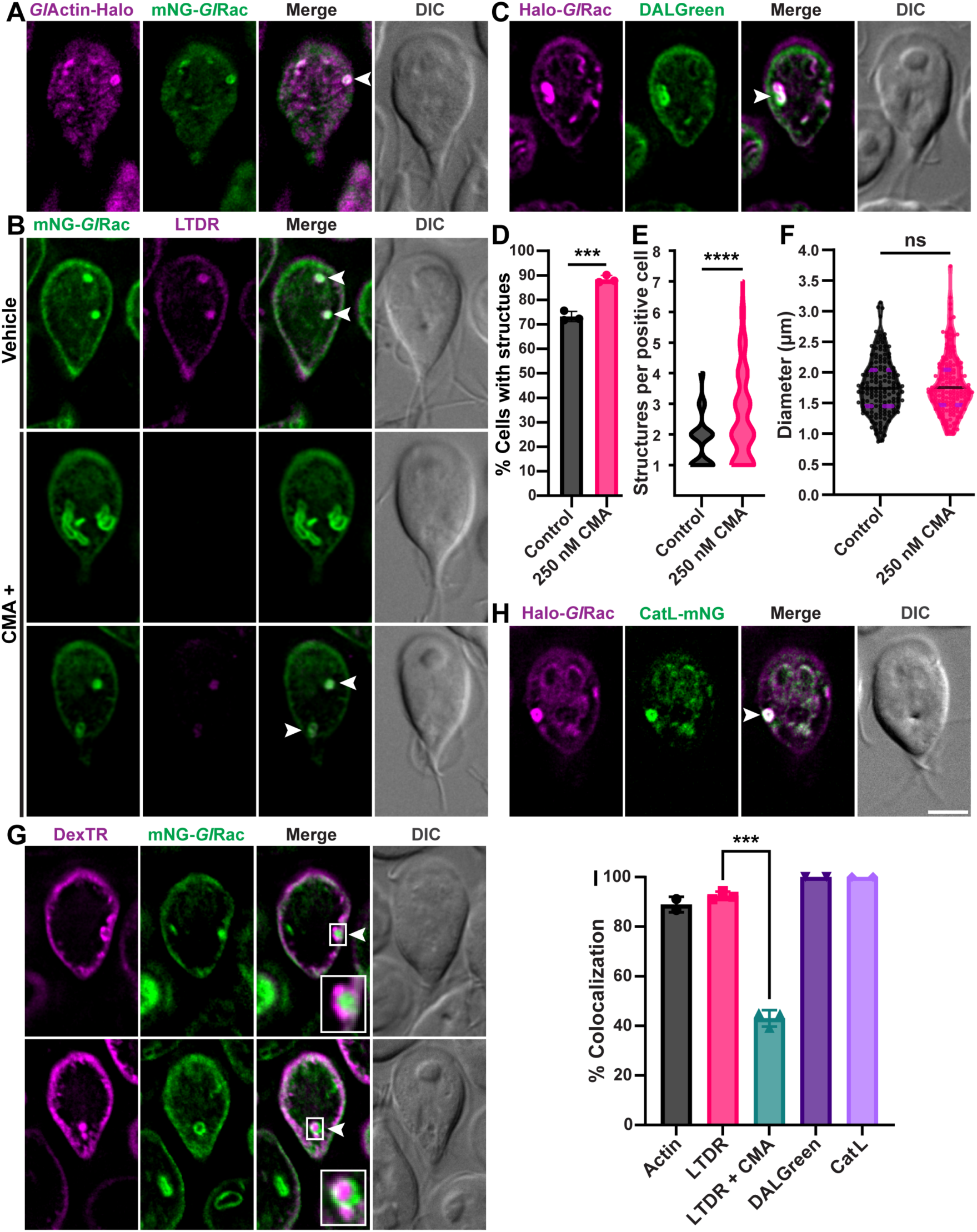
*Gl*Rac-positive compartments exhibit conserved autophagy-associated features. The indicated strains were cultured for 72 h. **(A)** Cellular distribution of *Gl*Actin-Halo (GL50803_0040817-Halo; magenta) in the mNG-*Gl*Rac (green) cells. **(B)** Cellular distribution of LysoTracker Deep Red (LTDR; magenta) in control (top; acetonitrile vehicle) and concanamycin A-treated (CMA+; middle and bottom) mNG-*Gl*Rac cells. Representative examples illustrate both the loss and retention of LTDR labeling following CMA treatment. **(C)** Cellular distribution of DALGreen (green) in the Halo-*Gl*Rac (magenta) strain. **(D)** Quantification of cells containing mNG-*Gl*Rac-labeled structures in control and CMA-treated cells. Trophozoites were treated with 250 nM CMA or vehicle during the final 24 h of culture. For each condition, n > 500 cells. Data represent mean ± SD from three independent experiments. P-values were calculated using paired two-tailed t-tests (***P ≤ 0.001). **(E)** Violin plot showing the distribution of the number of *Gl*Rac-positive structures in positive cells (cells containing at least one structure) in control and 250 nM CMA-treated cultures (n = 105 cells from three independent experiments). P-values were calculated using two-tailed Welch’s t-tests (****P ≤ 0.0001). **(F)** Violin plot showing the size distribution (maximum diameter) of spherical compartments measured in control and 250 nM CMA-treated cultures (all compartments from n = 100 cells, three independent experiments). Each dot represents one measurement. The solid black line marks the median, and the dashed purple lines indicate the first and third quartiles. P-values were calculated using two-tailed Welch’s t-tests (ns, not significant, P > 0.05). **(G)** mNG-*Gl*Rac cells treated with 1 mg/mL Dextran Texas Red (DexTR; magenta) for 24 h exhibit polysaccharide accumulation in peripheral vacuoles surrounding (top panel) and inside (bottom panel) *Gl*Rac-positive organelles. Insets show magnified views of the organelles. DexTR images are representative of at least 10 similar events across three independent experiments. **(H)** Cellular distribution of cathepsin L (GL50803_009548-mNG, green) in the Halo-*Gl*Rac strain. White arrowheads indicate colocalization with *Gl*Rac-positive structures. Scale bar, 5 µm. **(I)** Quantification of cells showing colocalization between *Gl*Rac-labeled structures and each element shown in (A-C) and (H). “P-values were calculated using paired two-tailed t-tests (***P ≤ 0.001). For each condition, n > 200 cells from at least two independent experiments.

A conserved feature of autophagosome maturation is compartment acidification following fusion with lysosomes in mammals, or with the vacuole in yeast and plants, enabling cargo degradation and nutrient recycling (1,9,63). To assess whether *Giardia*’s *Gl*Rac-positive compartments are acidic, we stained nutrient-depleted mNG-*Gl*Rac cells with LysoTracker Deep Red, a lysosomotropic dye that accumulates in low-pH organelles (36,40). This probe labeled 92.5% of *Gl*Rac-positive structures (Fig 5B and I), indicating that they are acidified. As expected, LysoTracker also stained PVs, which serve as acidic vacuoles in *Giardia* (22). Similar results were obtained using DALGreen, a pH-sensitive live-cell autophagy probe (79), which labeled both PVs and, more intensely, all *Gl*Rac-positive structures in Halo-*Gl*Rac cells (Fig 5C and I).

To determine whether V-ATPase activity is required for the acidification of *Gl*Rac-positive compartments, we treated trophozoites with concanamycin A, a specific V-ATPase inhibitor (80). As expected, concanamycin A markedly reduced LysoTracker labeling (Fig. 5B). However, 43% of *Gl*Rac-positive compartments retained detectable signal (Fig. 5B and I), indicating that a subset of these compartments remained partially acidified following V-ATPase inhibition. Concanamycin A treatment also increased the proportion of cells containing Halo-*Gl*Rac-labeled structures from 73.2% in control cultures to 88.4% following treatment (Fig. 5D). Concanamycin A shifted the distribution of *Gl*Rac-positive structures toward higher numbers per cell, decreasing the proportion of cells containing one or two structures while increasing the frequency of cells containing three to seven structures (Fig 5E). Although the average diameter of individual spherical compartments was not significantly altered by concanamycin A treatment (control: 1.77 µm, range 0.87–3.14 µm; CMA: 1.78 µm, range 0.99–3.74 µm; Fig 5F), the frequency of closely apposed, beaded clusters (Fig. 5B) increased from 17.2% in control cultures to 46.3% following treatment, replacing the predominance of discrete spherical structures.

Since *Giardia* lacks canonical lysosomes, we hypothesized that PVs, the parasite’s hybrid endosome/lysosome-like vacuoles, contribute to the acidification of *Gl*Rac-positive organelles. Supporting this idea, TEM analysis revealed PVs surrounding these organelles (Fig 1C and S2 Fig). To test this hypothesis, we incubated mNG-*Gl*Rac cells with Dextran Texas Red 10,000 MW (DexTR), a fluid-phase marker taken up in bulk by PVs (81,82), during the final 24 h of a 72 h culture. *Gl*Rac-labeled compartments were frequently observed adjacent to DexTR-labeled PVs, and in some cases, the DexTR signal was also detected within these organelles (Fig 5G). However, association with DexTR-positive material was heterogeneous, and some compartments displayed little or no detectable DexTR signal (S9 Fig). Together, these observations indicate that *Gl*Rac-labeled organelles can interact with DexTR-labeled PVs, consistent with the hypothesis that PV fusion could contribute to organelle acidification.

Cathepsins, a family of acidic proteases, mediate the degradation of autophagosomal contents, thus representing another key component in the autophagic response (83). To identify cathepsins potentially involved in *Giardia’s* autophagic-like response, we analyzed existing RNA-seq data profiling trophozoite transcriptomes during log (48 h) and declining (96 h) growth phases (84). We selected seven cathepsins that were upregulated by at least 2-fold at 96 h compared with 48 h (S1 Dataset), generated C-terminal mNG fusions, and expressed them in the Halo-*Gl*Rac background. Under nutrient-depleted conditions, four of these cathepsins were detectable and were consistently recruited to Halo-*Gl*Rac-labeled structures. The most upregulated cathepsin, based on RNA sequencing data, is shown in Fig 5H, while the additional cathepsins are presented in S10 Fig.

### *Gl*Rac-positive organelles harbor active cysteine proteases

Although several cathepsins localized to *Gl*Rac-positive compartments, localization alone does not demonstrate proteolytic activity. We therefore used the activity-based probe BODIPY-LHVS, which covalently labels active cysteine proteases while preserving their subcellular localization (85). BODIPY-LHVS labeling revealed active cysteine proteases in *Gl*Rac-positive organelles, with labeling also observed in PVs and the ER (Fig 6A), as previously reported (81). Pretreatment with the unlabeled inhibitor LHVS markedly reduced BODIPY-LHVS labeling (Fig 6B), confirming that probe accumulation reflects specific labeling of active cysteine proteases rather than nonspecific accumulation within these compartments. Together, these observations demonstrate that *Gl*Rac-positive organelles contain active cysteine proteases.

**Fig 6.**
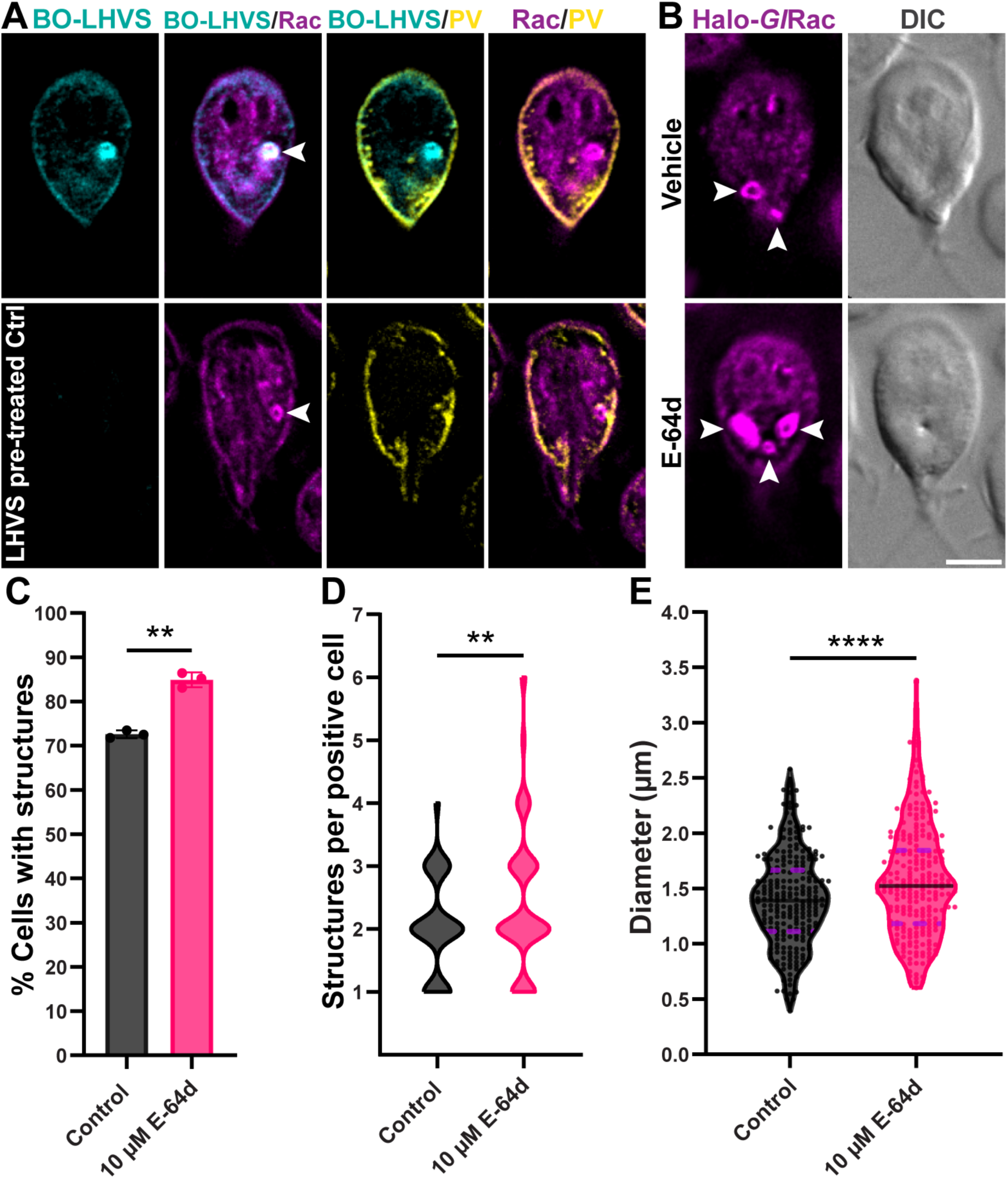
*Gl*Rac-positive organelles contain active cysteine proteases and increase in abundance and size following E-64d treatment. **(A)** Representative images of Halo-*Gl*Rac (magenta) cells labeled with the activity-based probe BODIPY-LHVS (BO-LHVS; cyan; top panel), which covalently labels active cysteine proteases. Cells were simultaneously incubated with DexTR to label peripheral vacuoles (PV; yellow). BO-LHVS labeling was detected throughout the endoplasmic reticulum but was enriched in PVs and *Gl*Rac-positive organelles (white arrowhead). Pretreatment with unlabeled LHVS markedly reduced BO-LHVS labeling (bottom panel), confirming the specificity of probe labeling. BO-LHVS and LHVS-pretreated control images were acquired using the same exposure settings and processed with identical intensity scaling and are representative of at least 10 similar events across two independent experiments. Scale bar, 5 µm. **(B)** Representative images of live Halo-*Gl*Rac cells cultured for 72 h and treated with 10 µM E-64d or DMSO (vehicle) during the final 24 h. Scale bar, 5 µm. **(C)** Quantification of cells containing Halo-*Gl*Rac-labeled structures in control and E-64d-treated cells. For each condition, n > 450 cells. Data represent mean ± SD from three independent experiments. P-values were calculated using paired two-tailed t-tests (**P ≤ 0.01). **(D)** Violin plot showing the distribution of the number of *Gl*Rac-positive structures in positive cells (cells containing at least one structure) in control and 10 µM E-64d-treated cultures (n = 105 cells from three independent experiments). P-values were calculated using two-tailed Welch’s t-tests (**P ≤ 0.01). **(E)** Violin plot showing the size distribution (maximum diameter) of spherical compartments measured in control and 10 µM E-64d-treated cultures (all compartments from n = 100 cells, three independent experiments). Each dot represents one measurement. The solid black line marks the median, and the dashed purple lines indicate the first and third quartiles. P-values were calculated using two-tailed Welch’s t-tests (****P ≤ 0.0001).

Because cathepsins B, L, and L-like were robustly recruited to Halo-*Gl*Rac-labeled structures, we used E-64d, a membrane-permeable inhibitor of cathepsins B, H, and L (86,87), to test whether cysteine protease inhibition affects the abundance or morphology of these structures under nutrient-depleted conditions. E-64d treatment increased the proportion of cells containing Halo-*Gl*Rac–labeled structures from 72.6% in controls to 84.9% (Fig 6C). While most positive cells in both conditions contained two to three compartments, E-64d broadened the distribution from 1-4 per cell in controls to 1-6, increasing the frequency of cells with higher compartment numbers (Fig 6D). E-64d also increased organelle size, with mean diameter expanding from 1.40 µm (0.39–2.58 µm) to 1.57 µm (0.60–3.38 µm; Fig 6E). Together, these findings indicate that cysteine protease activity contributes to the turnover of *Gl*Rac-positive organelles and that inhibition of this activity results in their accumulation and enlargement.

### *Gl*Rac regulates degradative organelle formation

Rho GTPases are well-established regulators of autophagy in other eukaryotes, including *Arabidopsis* and mammalian cells (40,41). Since *Gl*Rac is the sole Rho GTPase in *Giardia* and localizes to organelles exhibiting conserved autophagy-associated features, we investigated whether it affects the proportion of cells harboring these structures using both loss- and gain-of-function approaches.

To assess loss-of-function, we used a validated translation-blocking morpholino that reduces *Gl*Rac levels by approximately 70% at 24 h post-transfection (39,88). Depletion was confirmed by Western blotting (S11 Fig) and by reduced Halo-*Gl*Rac signal in fluorescence microscopy (Fig 7A and S12A Fig). Following *Gl*Rac depletion and exposure to nutrient-depleted medium, the proportion of cells containing *Gl*Rac-positive compartments decreased by 39% relative to the control (Fig 7B).

**Fig 7.**
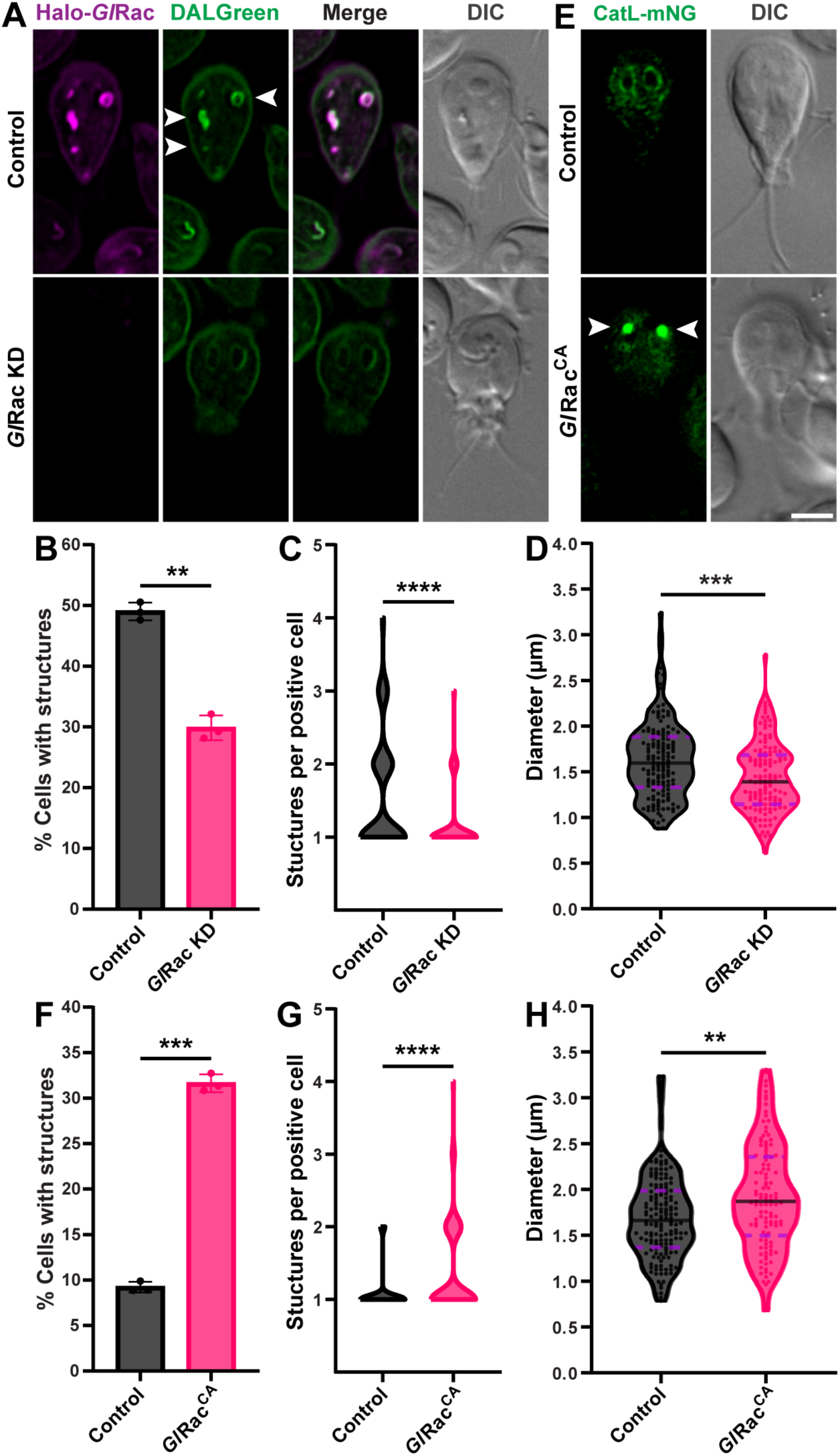
*Gl*Rac bidirectionally regulates degradative compartment formation. **(A)** Representative images of Halo-*Gl*Rac^MS^ cells 24 h after transfection with either a nonspecific control morpholino (control) or a translation-blocking morpholino targeting *Gl*Rac (*Gl*Rac KD). White arrowheads indicate *Gl*Rac-positive structures. **(B)** Quantification of morpholino-sensitive Halo-*Gl*Rac (Halo-*Gl*Rac^MS^) cells containing *Gl*Rac-positive structures after transfection with either a translation-blocking morpholino targeting *Gl*Rac (*Gl*Rac KD) or a nonspecific control (control). Between 18.5 and 24 h post-electroporation, cultures were exposed to nutrient-depleted medium from 72 h cultures to induce the formation of *Gl*Rac-positive structures, which were labeled with DALGreen. For each condition, n > 260 cells. Data represent mean ± SD from three independent experiments. P-values were calculated using paired two-tailed t-tests (**P ≤ 0.01). **(C)** Violin plot showing the distribution of the number of *Gl*Rac-positive structures in positive cells (cells containing at least one structure) in control and *Gl*Rac KD conditions (n = 105 cells from three independent experiments). P-values were calculated using two-tailed Welch’s t-tests (****P ≤ 0.0001). **(D)** Violin plot showing the size distribution (maximum diameter) of spherical compartments measured in control and *Gl*Rac KD conditions (all compartments from n = 100 cells across three independent experiments). Each dot represents one measurement. The solid black line marks the median, and the dashed purple lines indicate the first and third quartiles. P-values were calculated using two-tailed Welch’s t-tests (***P ≤ 0.001). **(E)** Representative images of cells expressing constitutively active *Gl*Rac (*Gl*Rac^CA^) or Halo-*Gl*Rac (control) after tetracycline treatment for 24 h in log-phase cultures. White arrowheads indicate *Gl*Rac-positive structures. Scale bar, 5 µm. **(F)** Quantification of cells containing degradative structures after tetracycline-induced expression of *Gl*Rac^CA^ for 24 h in log-phase growth cultures. Halo-*Gl*Rac cells (control) were grown under the same conditions. *Gl*Rac-positive structures were labeled with the cathepsin L-like (CatL) protease GL50803_00137680-mNG, which coincides with Halo-*Gl*Rac-labeled compartments (S10 Fig). For each condition, n > 300 cells. Data represent mean ± SD from three independent experiments. P-values were calculated using paired two-tailed t-tests (***P ≤ 0.001). **(G)** Violin plot showing the distribution of the number of *Gl*Rac-positive structures in positive cells (cells containing at least one structure) in the control and *Gl*Rac^CA^ (n = 105 cells from three independent experiments). P-values were calculated using two-tailed Welch’s t-tests (****P ≤ 0.0001). **(H)** Violin plot showing the size distribution (maximum diameter) of spherical compartments measured in the control and *Gl*Rac^CA^ (all compartments from n = 100 cells across three independent experiments). Each dot represents one measurement. The solid black line marks the median, and the dashed purple lines indicate the first and third quartiles. P-values were calculated using two-tailed Welch’s t-tests (**P ≤ 0.01).

Within the *Gl*Rac morpholino-treated population, a subset of cells retained high Halo-*Gl*Rac signal and normal morphology (“bright” cells), while others showed reduced signal and severe morphological defects (“dim” cells), as previously reported for *Gl*Rac-depleted cells (39) (S12A Fig). Our initial quantification focused on dim cells, which exhibit stronger depletion and typically more pronounced defects. However, even when the entire population (both bright and dim cells) was considered, the proportion of cells containing *Gl*Rac-positive structures remained significantly lower than in the control (S12B Fig).

To determine whether *Gl*Rac depletion affects not only the frequency but also the properties of these degradative compartments, we quantified the number and size of compartments in dim *Gl*Rac KD cells. In the control, positive cells commonly contained multiple compartments (range: 1-4), and cells with two or more compartments were frequent (Fig 7C). In contrast, *Gl*Rac-depleted cells showed a marked shift toward fewer compartments per cell, with almost all positive cells containing only a single compartment (range: 1-3) (Fig 7C). Thus, *Gl*Rac loss reduces the likelihood that a cell will form multiple degradative compartments. *Gl*Rac depletion also affected organelle size. Spherical compartments decreased from an average diameter of 1.63 µm in controls (range: 0.88-3.24 µm) to 1.44 µm in *Gl*Rac KD cells (range: 0.61-2.78 µm) (Fig 7D).

To assess gain of function, we used a constitutively active GTP-locked *Gl*Rac mutant (tetracycline-inducible Q74L HA-*Gl*Rac; HA-*Gl*Rac^CA^) (Fig 7E) previously shown to trigger the formation of large vesicular structures in non-encysting trophozoites, indicating a role for *Gl*Rac in endomembrane organization (38,39). Because gain-of-function and loss-of-function experiments require opposite baselines, we assayed *Gl*Rac^CA^ in log-phase cultures, where *Gl*Rac-positive compartments are scarce, rather than the compartment-rich conditions used for knockdown. Induction of HA-*Gl*Rac^CA^ expression for 24 h increased the proportion of cells harboring *Gl*Rac-positive structures from 9.2% in uninduced controls to 31.6% (Fig 7F).

We next quantified the number and size of *Gl*Rac-positive compartments in positive cells. In control cultures, most positive cells contained a single compartment, with a narrow distribution ranging from 1-2 compartments per cell (Fig 7G). In contrast, *Gl*Rac^CA^ expression broadened this distribution to 1-4 compartments per cell, with a pronounced increase in cells containing multiple compartments (Fig 7G). Thus, constitutive activation of *Gl*Rac enhances not only the frequency of cells that form *Gl*Rac-positive structures but also the number of compartments per cell. *Gl*Rac activation also increased organelle size. Spherical *Gl*Rac-positive compartments in control cells averaged 1.70 µm in diameter (range: 0.79-3.23 µm), whereas those in *Gl*Rac^CA^-expressing cells increased to an average of 1.92 µm (range: 0.68-3.31 µm) (Fig 7H).

Together, these findings demonstrate that *Gl*Rac acts as a bidirectional regulator of *Gl*Rac-positive compartment biogenesis in *Giardia*: its depletion reduces compartment formation, whereas its constitutive activation enhances it. This behavior is consistent with roles reported for Rho GTPases in autophagy initiation and membrane remodeling in other eukaryotic systems.

### Quinacrine treatment promotes the accumulation of *Gl*Rac-positive structures

Quinacrine, originally developed as an antimalarial drug, is also used to treat nitroimidazole-refractory giardiasis, although its mechanism of action in *Giardia* remains unclear (89–91). Notably, this compound concentrates in acidic organelles and inhibits autophagic flux in mammalian cells, leading to the accumulation of autophagosomes (92,93). Intriguingly, quinacrine treatment has been reported to induce dense cytoplasmic blebs in *Giardia*, resembling the *Gl*Rac-positive organelles described in this study (94).

To investigate how quinacrine affects *Gl*Rac-positive structures, Halo-*Gl*Rac cells were treated with 10 µM quinacrine for 24 h. Taking advantage of the drug’s green autofluorescence, we observed its accumulation in PVs and within or surrounding *Gl*Rac-positive structures (Fig 8A and B). This pattern is consistent with compartment acidification, as quinacrine concentrates in low-pH organelles via an ion-trapping mechanism (95,96).

**Fig 8.**
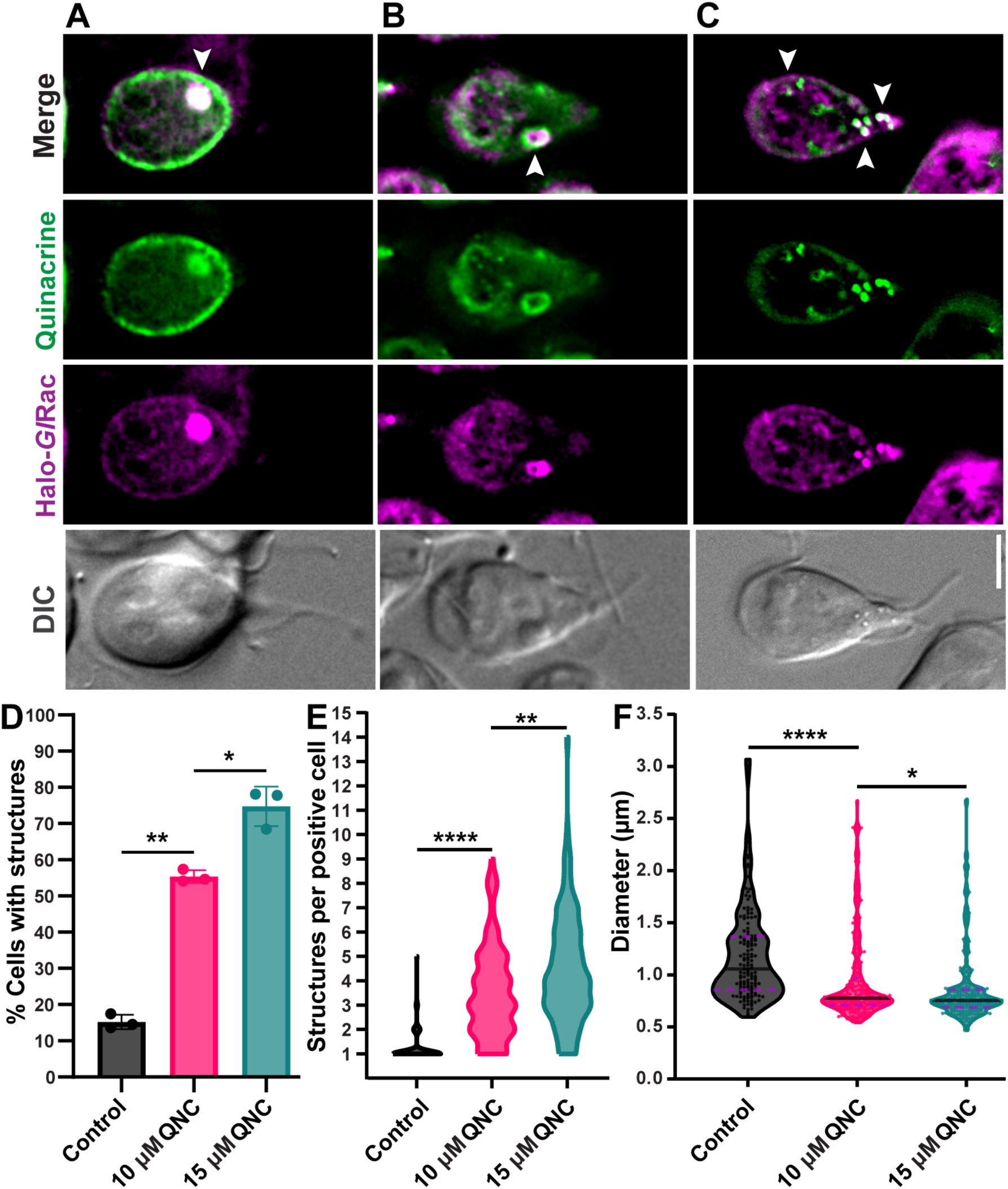
Quinacrine treatment promotes the accumulation of *Gl*Rac-positive structures. **(A-C)** Representative images of live Halo-*Gl*Rac cells treated with 10 µM quinacrine for 24 h. Quinacrine (green autofluorescence) accumulates in peripheral vacuoles and Halo-*Gl*Rac-labeled compartments **(A)** and is also detected surrounding these compartments **(B)**. Quinacrine treatment additionally induces small, intensely fluorescent structures that overlap with Halo-*Gl*Rac signal **(C)**; these *Gl*Rac-positive structures were included in the quantification. White arrowheads indicate the features described in each panel. Scale bar, 5 µm. **(D)** Quantification of Halo-*Gl*Rac cells harboring *Gl*Rac-positive structures after being cultured for 24 h, followed by treatment with vehicle (Control), 10 µM, or 15 µM quinacrine (QNC) for 24 h. For each condition, n > 300 cells. Data are presented as mean ± SD from three independent experiments. P-values were calculated using paired two-tailed t-tests (*P ≤ 0.05; **P ≤ 0.01). **(E)** Violin plot showing the distribution of the number of *Gl*Rac-positive structures in positive cells (cells containing at least one structure) in the control, 10 µM, and 15 µM quinacrine treatments (n = 105 cells from three independent experiments). P-values were calculated using two-tailed Welch’s t-tests (**P ≤ 0.01; ****P ≤ 0.0001). **(F)** Violin plot showing the size distribution (maximum diameter) of spherical Halo-*Gl*Rac-labeled compartments measured in control, 10 µM, and 15 µM quinacrine treatments (all compartments from n = 100 cells across three independent experiments). Each dot represents one measurement. The solid black line marks the median, and the dashed purple lines indicate the first and third quartiles. P-values were calculated using two-tailed Welch’s t-tests (*P ≤ 0.05; ****P ≤ 0.0001).

In addition to the compartments previously described, quinacrine treatment induced smaller, intensely fluorescent structures that frequently colocalized with Halo-*Gl*Rac (Fig 8C). While it remains unclear whether these structures are equivalent to the *Gl*Rac-positive compartments observed during nutrient depletion or encystation, their consistent and intense Halo-*Gl*Rac labeling suggests a functional relationship. Similar phenotypes have been reported in mammalian cells, where quinacrine induces the formation of immature or defective autophagolysosomes, often smaller than canonical autophagosomes (93,97). We therefore included these *Gl*Rac-labeled structures in our quantification. Under these conditions, quinacrine treatment markedly increased the proportion of cells containing *Gl*Rac-positive structures, from 15.2% in the control to 55.4% when treated with 10 µM quinacrine (Fig 8D). A higher concentration (15 µM) further raised this proportion to 74.8%, demonstrating a dose-dependent effect (Fig 8D).

Furthermore, in the control, positive cells typically contained a single compartment (range: 1-5) (Fig 8E). Quinacrine treatment substantially broadened this distribution: at both 10 µM and 15 µM, the number of compartments per cell ranged from 1-9 and 1-14, respectively, with three compartments becoming the most common outcome (Fig 8E). Thus, quinacrine promotes a dose-dependent accumulation of *Gl*Rac-positive compartments. Quinacrine also affected organelle size (Fig 8F). Spherical *Gl*Rac-positive organelles decreased in average diameter from 1.18 µm in the control (range: 0.60-3.07 µm) to 0.95 µm at 10 µM (range: 0.54-2.67 µm) and 0.86 µm at 15 µM (range: 0.47-2.69 µm) (Fig 8F), a shift consistent with the accumulation of smaller, immature compartments often observed under autophagic flux inhibition caused by this drug.

Together, these observations indicate that quinacrine disrupts the autophagic-like response in *Giardia*, increasing both the number of *Gl*Rac-positive compartments and the proportion of cells that form them, paralleling its effects in other eukaryotic systems.

## Discussion

Autophagy is widely recognized as an ancient and essential eukaryotic process, yet its presence in *Giardia* has been questioned because the parasite lacks canonical lysosomes and several conserved ATG proteins, most notably ATG8 (5,6,10,11,19). Here we show that *Gl*Rac marks a distinct class of degradative compartments with multiple properties of autophagosomes in model eukaryotes. Their formation is induced by nutrient starvation and encystation and is negatively regulated by GTOR, consistent with the conserved role of TOR in coupling nutrient availability to autophagy induction. Morphologically, these compartments are bounded by double and multilamellar membranes, a hallmark of autophagosomes. Functionally, they acidify, label with the autolysosome reporter DALGreen, and recruit active cathepsin proteases. Perturbing degradative turnover, either by inhibiting cysteine proteases with E-64d or by disrupting acidification with the V-ATPase inhibitor concanamycin A, caused *Gl*Rac-positive compartments to accumulate, although the resulting morphologies differed between the two treatments. E-64d promoted the accumulation of enlarged individual compartments, whereas concanamycin A increased the frequency of closely apposed, beaded compartment clusters, as expected for structures that are continuously formed, matured, and cleared rather than static end points. Their biogenesis requires *Gl*Rac activity and is accompanied by robust actin recruitment, consistent with the conserved role of actin in autophagosome formation (77,98). Together, these features identify a degradative pathway that is functionally analogous to autophagy (Fig 9). Notably, however, this pathway relies on a markedly reduced complement of ATG-like factors and forms over a substantially slower timescale than autophagosomes in yeast and mammalian cells. We therefore conclude that *Giardia* possesses a highly divergent, ATG8-independent autophagy-like pathway in which *Gl*Rac serves as a central regulator of compartment formation (Fig 9).

**Fig 9.**
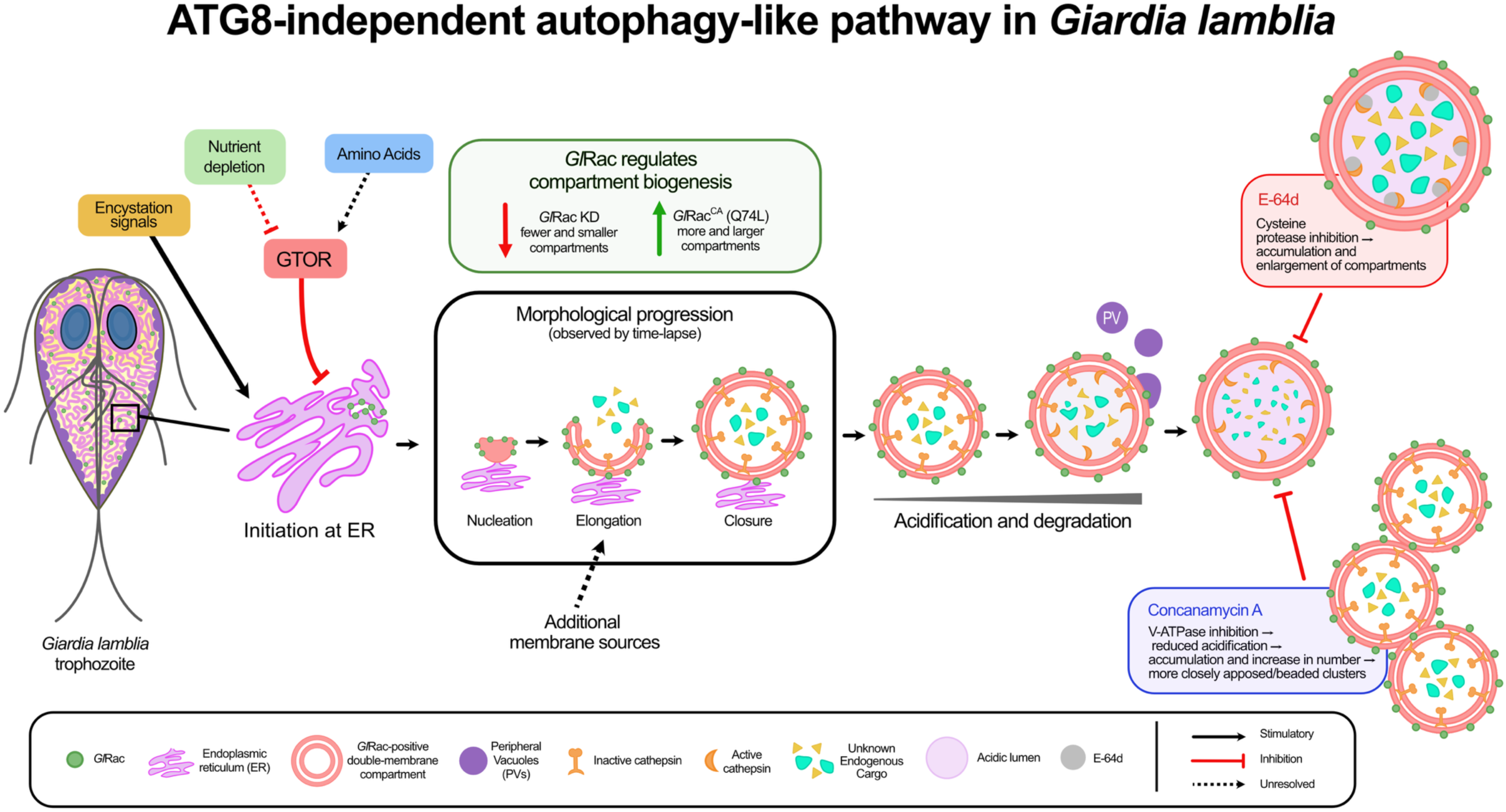
Working model for an ATG8-independent autophagy-like pathway in *Giardia lamblia*. Nutrient depletion and encystation promote the formation of *Gl*Rac-positive double-membrane compartments that initiate predominantly at the endoplasmic reticulum (ER). Compartment formation is negatively regulated by GTOR, whose activity is amino acid–responsive. Live-cell imaging revealed a morphological progression from nucleation through elongation and closure. *Gl*Rac regulates compartment biogenesis bidirectionally, as constitutive activation (*Gl*Rac^CA^) increases compartment abundance and size of compartments, whereas *Gl*Rac knockdown (*Gl*Rac KD) reduces them. *Gl*Rac-positive compartments can associate with peripheral vacuoles (PVs), although the mechanism and functional significance of these interactions remain unresolved. The compartments acquire degradation-associated features, including acidification and active cysteine proteases. Cysteine protease inhibition with E-64d promotes compartment accumulation and enlargement, whereas inhibition of V-ATPase-mediated acidification with concanamycin A promotes accumulation and an increase in compartment number, both consistent with impaired degradative turnover. Dashed arrows indicate proposed steps or components that remain unresolved, including additional membrane sources contributing to compartment biogenesis and the identity of endogenous cargo. For clarity, some unresolved features, including the upstream activator of *Gl*Rac and the effectors that recruit actin, are not depicted.

Previous live-cell GFP localization studies described unexplained inclusions or membrane-bound structures in *Giardia*, but these structures were not assigned to a defined organelle class or linked to a specific cellular pathway (37). The morphology, size, and prevalence of the *Gl*Rac-positive compartments described here raise the possibility that a subset of those structures corresponds to this autophagy-like pathway.

The pathway is tightly linked to nutrient availability. The proportion of cells harboring *Gl*Rac-positive compartments increased in older, nutrient-depleted cultures and declined rapidly after amino acid replenishment. Additionally, nutrient limitation increased the number of compartments per cell and modestly expanded their diameter. Consistent with conserved TOR-dependent control of autophagy (6,58,59) and with the amino acid–responsive autophagy described in other protozoan parasites such as *T. brucei*, *T. gondii*, and *P. falciparum* (53–56), GTOR expression decreased during starvation, and knockdown of GTOR with translation-blocking morpholinos increased compartment abundance, identifying GTOR as a negative regulator of compartment formation. Rapamycin did not phenocopy this effect, possibly because the FKBP–rapamycin binding domain is poorly conserved in *Giardia* (<30% identity) (60), reducing drug sensitivity.

Beyond nutrient status, this pathway is developmentally regulated: *Gl*Rac-positive compartments also form during encystation, the differentiation program that converts the replicative trophozoite into the environmentally resistant, transmissible cyst. This mirrors a recurring theme in protozoan parasites, in which autophagy is induced during life-cycle transitions and supports the morphological remodeling and metabolic adaptation those transitions demand (6,46–48,99). Encystation is a particularly remodeling-intensive transition, requiring cytoskeletal reorganization, flagellar retraction, ventral disc disassembly, and nuclear division, accompanied by a metabolic switch in which glycolytic carbon is diverted toward UDP-GalNAc synthesis to build the GalNAc homopolymer of the cyst wall, increasing the cell’s reliance on the arginine dihydrolase pathway for ATP (100–103). A regulated degradative pathway that mobilizes cellular material and membrane during this reorganization would be well suited to meet these demands. Because the cyst is the transmissible form of the parasite, a degradative pathway engaged during its formation may also bear on transmission and persistence.

These findings challenge the longstanding assumption that *Giardia* lacks autophagy and instead support the idea that this pathway has been adapted to operate with a minimal set of components, reflecting its parasitic lifestyle and reduced cellular complexity. Although *Giardia* exhibits several features consistent with an autophagy-like degradative pathway, the apparent absence of several key ATG proteins, particularly ATG8 (5,11,36,104), raises important questions about how this process is executed in this organism. One possibility is that *Giardia* encodes a highly divergent repertoire of ATG proteins undetectable by standard sequence-based tools. Similar cases have been described in other protists, where functionally conserved ATGs show low sequence identity to known counterparts (5,6). For example, the ATG12 conjugation system was initially thought to be absent in trypanosomatids (105). However, in *L. major*, ATG5, ATG10, and a protein annotated as ATG12 could functionally complement *S. cerevisiae* knockouts, despite their limited sequence similarity (47,106). Furthermore, in apicomplexan parasites such as *Toxoplasma sp.* and *Plasmodium spp.*, ATG12 and ATG5 form a non-covalent complex that bypasses the E2-like enzyme ATG10, while still enabling ATG8 lipidation (107). These examples highlight that the absence of a recognizable ortholog does not necessarily equate to loss of pathway function. Divergent protist lineages may instead rely on alternative proteins or mechanisms to perform roles characterized initially in model organisms. In some cases, functionally equivalent proteins evolve independently or are replaced by unrelated or broadly acting components (5,6,108).

Another non-mutually exclusive possibility is that *Giardia* performs a highly reduced form of autophagy that relies only on the most essential components of the pathway. Although ATG8 and its conjugation machinery have long been considered essential, growing evidence from mammalian systems demonstrates that autophagosome formation can still occur in their absence (109–112). For instance, cells lacking all six mammalian ATG8 paralogs still form autophagosomes, albeit with reduced efficiency (109,112), and recent genetic screens further show that some cargos can be degraded independently of ATG8 lipidation (113). In *Giardia*, the relatively slow appearance of autophagosome-like compartments, ranging from 30 min to 8 h, may likewise reflect reduced efficiency in the absence of ATG8, compared to the rapid formation (≤10 min) typically observed in yeast and mammalian cells upon autophagy induction (66–68). Together, these findings highlight the evolutionary plasticity of the autophagy machinery and suggest that ATG8-independent mechanisms may be more common than previously anticipated.

The first experimental localization of the candidate ATG proteins in *Giardia* did not identify a robust compartment marker comparable to *Gl*Rac. Although some candidates showed coincident signal at these compartments, most remained broadly distributed rather than enriched. Because many ATGs act only transiently or at specific stages, stage-restricted recruitment cannot be excluded. Thus, localization is limited both spatially and temporally, and cannot on its own establish functional involvement. Assigning ATG-like roles to these candidates will require direct functional tests, such as knockdown or overexpression.

Considering the functional and localization data together with reciprocal BLAST results, we grouped these candidates into four categories (S1 Dataset). GTOR is supported by both sequence and functional evidence: it recovers TOR on reciprocal search, and its depletion increases compartment abundance. *Gl*ATG1.1, *Gl*VPS15, *Gl*VPS34, and *Gl*ATG21 recover the expected protein on reciprocal search but remain functionally untested, and we regard them as putative. *Gl*ATG1.2 and *Gl*ATG18 recover a related family member rather than the expected ortholog, indicating that family membership is supported, whereas the specific ortholog assignment remains uncertain. For three candidates, the two lines of evidence converge against the prior assignment. GL50803_0033762, proposed as ATG16, recovers POC1 at 2 × 10^−83^ and localizes to cytoplasmic axonemes. GL50803_006288 and GL50803_0028994, proposed as ATG7 and ATG8, return no significant hit at default settings and localize to nuclei and to only a small minority of *Gl*Rac-positive compartments, respectively; AlphaFold does not predict a ubiquitin fold for the latter. We therefore withdraw these three assignments.

Rho GTPases are central regulators of the actin cytoskeleton, and this role is evident in *Giardia* (38). Actin, a conserved mediator of autophagosome dynamics, is robustly recruited to *Gl*Rac-labeled compartments, implicating cytoskeletal remodeling in their formation. How *Gl*Rac promotes this recruitment remains unclear. In other eukaryotes, Rho-family GTPases drive local actin assembly by activating nucleation-promoting factors that stimulate the Arp2/3 complex (114,115). *Giardia*, however, lacks recognizable components of this pathway, including the Arp2/3 complex, its nucleation-promoting factors, and the canonical actin-binding proteins that would otherwise couple *Gl*Rac to *Gl*Actin (116). Thus, *Gl*Rac likely acts through divergent or unidentified effectors.

Cytoskeletal remodeling is unlikely to be *Gl*Rac’s only contribution to compartment biogenesis. *Gl*Rac associates with membranes and interacts with several Rab GTPases (117), consistent with its established role in *Giardia* membrane trafficking (38,39). The *Gl*KDELR labeled 81.5% of *Gl*Rac-positive compartments and closely colocalized with *Gl*Rac, consistent with the ER association previously observed for *Gl*Rac by antibody staining (39). These findings identify the endoplasmic reticulum as a major membrane source for *Gl*Rac-positive compartments, paralleling the central role of the ER in autophagosome biogenesis in other eukaryotes (118–120). Because some compartments lacked detectable *Gl*KDELR signal, additional membrane sources, including encystation-specific secretory vesicles, mitosomes, or the plasma membrane, may also contribute.

The predominantly ER-derived character of these compartments also distinguishes them from *Giardia*’s PVs, the parasite’s endolysosomal system. Because PVs are frequently found in close apposition to *Gl*Rac-positive compartments (Figs 1C and 5G), one possible interpretation is that these structures represent a specialized subset of PVs. However, several observations argue against a PV origin. Ultrastructural analyses demonstrated that *Gl*Rac-positive compartments are bounded by double membranes and can develop multilamellar organization, features distinct from the single-membrane morphology of PVs. Time-lapse imaging revealed their de novo biogenesis through line-, cup-, and sphere-shaped intermediates, a developmental sequence distinct from the morphology of PVs. Finally, although DexTR was detected within a subset of *Gl*Rac-positive compartments, many others lacked detectable signal (S9 Fig), suggesting that interactions with the PV system are heterogeneous. Together, these observations place *Gl*Rac at ER-derived membranes during compartment formation and suggest that it coordinates both membrane and cytoskeletal remodeling. The compartments are not fully independent of the PV system, but they cannot be explained as nutrient depletion- or encystation-induced PVs.

In support of a functional degradative pathway, cathepsins B, L, and L-like are recruited to *Gl*Rac-positive compartments, and the activity-based probe BODIPY-LHVS detects active cysteine proteases at these compartments. Blocking these proteases with E-64d increased the proportion of cells containing compartments and promoted their accumulation and growth under nutrient-depleted conditions, altering both their abundance and morphology. In other eukaryotic systems, E-64d inhibits lysosomal cysteine proteases and reduces autophagic flux by blocking degradative turnover, leading to accumulation of autophagic compartments (36,86,87); the comparable response in *Giardia* indicates that proteolytic activity contributes to the turnover of these structures. Inhibiting compartment acidification with the V-ATPase inhibitor concanamycin A likewise promoted compartment accumulation but produced a distinct morphological phenotype, increasing the frequency of closely apposed, beaded compartment clusters rather than enlarging individual compartments. This altered morphology is consistent with impaired compartment maturation following V-ATPase inhibition. A similar accumulation of immature autophagic compartments has been reported after V-ATPase inhibition in model eukaryotes (121,122). Because acidification is required for cathepsin activation, E-64d and concanamycin A interfere with two sequential requirements of degradative turnover. Despite their distinct morphological outcomes, both perturbations resulted in compartment accumulation, supporting a model in which *Gl*Rac-positive compartments undergo continuous maturation and turnover rather than persisting as static structures.

Interestingly, in the context of identifying new therapeutic targets in *Giardia*, our findings show that quinacrine, a drug introduced for giardiasis treatment in the 1940s and currently used for nitroimidazole-refractory cases (90,91,94), accumulates in the acidic compartments, likely due to ion-trapping (95), and increases the proportion of cells harboring *Gl*Rac-positive compartments. In mammalian cells, this accumulation is driven by V-ATPase–dependent cation trapping and is accompanied by inhibition of autophagic flux, with reported buildup of degradation-incompetent autophagic compartments and compensatory lysosome biogenesis (92,93,97). That an established antigiardial drug engages this pathway is consistent with these compartments constituting a physiologically active degradative system and raises the possibility that the pathway contributes to the drug’s antigiardial activity (123). Quinacrine’s molecular targets in *Giardia* remain unknown, but these results connect a long-used antigiardial drug to a previously undescribed degradative pathway.

The divergence of parasite autophagy machinery from its mammalian counterparts (5,10,11) also offers a therapeutic opening: components distinct enough to be targeted selectively could yield parasite-specific inhibitors with minimal host toxicity. Consistent with this, a recent kinome-wide screen identified antigiardial compound clusters enriched for inhibitors of atypical protein kinases, including GTOR (124); although GTOR’s function remains incompletely defined, its presence among inhibitor-sensitive targets suggests that the nutrient-sensing and degradative signaling described here is pharmacologically vulnerable in *Giardia*. Together with the quinacrine data, these findings highlight this pathway as a promising source of therapeutic targets — an urgent need given the global burden of giardiasis, the limitations of current nitroimidazole therapy, and the demand for new antigiardial drugs (125–127).

Together, our findings establish *Gl*Rac as a central regulator of a divergent, ATG8-independent autophagy-like pathway in *Giardia*. This pathway integrates nutrient status, GTOR signaling, actin recruitment, acidification, and cathepsin-dependent turnover, and its induction during encystation suggests a role in differentiation-associated remodeling. The localization and effects of quinacrine further suggest that this previously unrecognized degradative pathway may be pharmacologically vulnerable. More broadly, *Giardia* demonstrates that the core logic of autophagy-like membrane remodeling can be preserved even when much of the canonical ATG machinery has been lost or radically altered.

## Materials and Methods

### Construct design

Coding sequences were PCR-amplified from *G. lamblia* genomic DNA using Q5 High-Fidelity DNA Polymerase (NEB) and primers listed in S2 Dataset, which also provides construct details. Parental NanoLuc, mNG and HaloTag vectors were digested with the appropriate restriction enzymes, and PCR products were assembled using the NEBuilder HiFi DNA Assembly Kit (NEB). Final constructs were verified by sequencing. Except for GL50803_003099-mNG and *Gl*Rac^CA^, all plasmids were linearized for integration using the restriction enzymes listed in S2 Dataset. DNA was purified using the DNA Clean & Concentrator-25 kit (Zymo Research) before being electroporated into *Giardia*. Linearized constructs were integrated into the endogenous locus by homologous recombination to generate endogenously tagged proteins (128). The integration strategy for the Halo-*Gl*Rac construct is shown in S1 Fig.

### *Giardia* strain and transfection

*G. lamblia* isolate WB clone C6 (ATCC 50803) was cultured at 37 °C in Keister’s modified TYI-S-33 medium (TYDK) adjusted to pH 7.1 and supplemented with 10% adult bovine serum and 0.52 mg/mL bovine bile (129). For transfections, 10-30 µg of purified plasmid was electroporated into 300 µL of chilled trophozoites (~8 × 10^6^ cells) using a 0.4 cm cuvette and a GenePulser XCell (Bio-Rad) set to 375 V, 1000 µF, and 750 Ω. Electroporated cells were transferred into 16 mL screw-cap round-bottom tubes (Corning Life Sciences) containing 13 mL of pre-warmed TYDK and incubated for 24 h at 37 °C before initiating antibiotic selection. G418 (InvivoGen; 115.4-307.7 µg/mL) or puromycin (InvivoGen; 11.5-30.8 µg/mL) concentrations were gradually increased for 7-14 days. During this selection period, the medium was replaced every 48 h.

### Modulation of *Gl*Rac-positive structures by encystation and nutrient availability

To induce encystation, the high bile method was used as previously described (130). Briefly, Halo-*Gl*Rac cells were incubated for 48 h in pre-encystation medium (PE), which is compositionally similar to TYDK but lacks bovine bile and is adjusted to pH 6.8. Cells were then transferred to encystation medium (TYI-S-33, pH 7.8) supplemented with 10% adult bovine serum, 10 g/L bovine and ovine bile (MilliporeSigma), and 5 mM calcium lactate. *Gl*Rac-positive structures were quantified at 0 (non-encysting cells in PE), 1, 2, 3, and 4 h post-encystation induction.

To induce nutrient depletion, Halo-*Gl*Rac cells were maintained in PE for 48, 72, or 96 h without medium replacement, and *Gl*Rac-positive structure levels were assessed at the end of the period. For refeeding experiments, 72 h cultures (0 h timepoint) were transferred to fresh TYDK and analyzed after 0.5, 1, 2, and 3 h. For individual nutrient supplementation, 10x stock solutions of glucose (100 g/L), casein digest (200 g/L), or L-arginine (100 mM) were prepared in TYDK salts (2 g/L NaCl, 0.6 g/L KH_2_PO_4_, 0.98 g/L K_2_HPO_4_, and 12 µg/mL ferric ammonium citrate). Cultures were supplemented with a single nutrient and analyzed after 2 h.

### GTOR morpholino knockdown

Morpholino knockdown experiments were performed as previously described (39,131). Briefly, Halo-*Gl*Rac cells were transfected with either a translation-blocking morpholino targeting GTOR (5′ AGTAATTGCGTCTGTGCTCATTTTT3′) or a standard control morpholino oligonucleotide (5′ CCTCTTACCTCAGTTACAATTTATA 3′) (Gene Tools LLC). After transfection, cells were incubated for 24 h in TYDK. GTOR depletion was confirmed by measuring the reduction in GTOR-NLuc signal using a plate reader (protocol described below).

### In vitro bioluminescence assays with GTOR

GTOR-NLuc cells were cultured in PE medium for 24 h or 72 h. After incubation, cells were placed on ice for 30 min and centrifuged at 700 × g for 7 min at 4 °C. Cell counting and plate reader assays followed published methods (130,132), with minor modifications. Pellets were resuspended in cold 1× HBS, and cell density was measured using a MOXI Z Mini Automated Cell Counter Kit (Orflo). For each reaction, 20,000 cells were dispensed into white, flat-bottom 96-well plates (Corning), mixed with 10 μL of NanoGlo Luciferase Assay Reagent (Promega) in a final reaction volume of 200 μL immediately before measurement, and assayed in triplicate. Relative luminescence units (RLU) were recorded on a pre-warmed SPARK plate reader (Tecan) at 37 °C for 30 min, and the maximum signal obtained during this period was used for analysis.

### Pharmacological treatments

To assess the effect of rapamycin, Halo-*Gl*Rac cells were cultured for 48 h in PE and then treated with 36 µM rapamycin (MilliporeSigma) for 2 h. DMSO-treated cells served as vehicle controls.

For concanamycin A treatment, mNG-*Gl*Rac cells were cultured for 48 h in TYDK and then treated with 250 nM concanamycin A (Cayman Chemical Co.) or vehicle (0.022% acetonitrile) for 24 h. The concentration of concanamycin A was selected based on preliminary dose-response experiments as the highest concentration that did not produce an obvious growth defect under the experimental conditions.

For E-64d treatment, Halo-*Gl*Rac cells were cultured for 48 h in PE and then treated with 10 µM E-64d (EST; Sigma) or vehicle (0.05% DMSO) for 24 h. The concentration of E-64d was selected based on previous reports showing effective inhibition of cysteine protease activity in *Giardia* trophozoites. Final DMSO concentrations were kept constant across conditions.

For quinacrine treatment, Halo-*Gl*Rac cells were cultured for 24 h in PE, then treated with 10 or 15 µM quinacrine dihydrochloride (MilliporeSigma) or vehicle (Milli-Q H_2_0) for 24 h. Quinacrine solutions were filtered using a Steriflip Vacuum Tube Top Filter (Millipore).

### Live-cell labeling of acidic and endocytic compartments

To assess compartment acidification, mNG-*Gl*Rac cells were cultured for 72 h, then incubated in HEPES-buffered saline (HBS; 137 mM NaCl, 5 mM KCl, 0.91 mM Na₂HPO₄·7H₂O, 5.55 mM glucose, 20 mM HEPES, pH 7.0) containing 75 nM LysoTracker Deep Red (Thermo Fisher Scientific) for 30 min before imaging. DALGreen labeling was performed on Halo-*Gl*Rac cells cultured for 72 h and incubated in HBS containing 0.2 µM DALGreen (Dojindo) for 30 min before imaging. For experiments evaluating the effect of concanamycin A on compartment acidification, mNG-*Gl*Rac cells were first treated with 250 nM concanamycin A or vehicle for the final 24 h of culture as described above, then incubated in HBS containing LysoTracker Deep Red.

To investigate the association between PVs and *Gl*Rac-positive compartments, mNG-*Gl*Rac cells were incubated for 48 h in PE, followed by incubation with 1 mg/mL Dextran Texas Red (10,000 MW, lysine-fixable; Invitrogen) for 24 h.

To visualize active cysteine proteases, Halo-*Gl*Rac cells from 72 h cultures in PE were incubated with 400 nM BODIPY-LHVS (85) for 1 h before imaging. For inhibition controls, cells were pretreated with 10 µM unlabeled LHVS cathepsin inhibitor (Millipore Sigma) for 1 h before BODIPY-LHVS labeling. During the final 1 h incubation, all samples were also incubated with 1 mg/mL Dextran Texas Red (10,000 MW, lysine-fixable; Invitrogen). Cells were imaged immediately using the live-cell imaging conditions described below.

### *Gl*Rac morpholino knockdown and constitutive activation

Morpholino knockdown experiments were performed as previously described (39,131). Briefly, morpholino-sensitive Halo-*Gl*Rac (Halo-*Gl*Rac^MS^) cells were transfected with either a translation-blocking morpholino targeting *Gl*Rac (5′ TATCCTCATTTCCTGTACTAGTCAT 3′) or a standard control morpholino oligonucleotide (5′ CCTCTTACCTCAGTTACAATTTATA 3′) (Gene Tools LLC). After transfection, cells were incubated for 18.5 h in TYDK, centrifuged at 700 × g for 7 min, and then transferred to nutrient-depleted PE from a 72 h culture for 5 h to induce the formation of *Gl*Rac-positive structures. *Gl*Rac depletion was evaluated by a reduction in Halo-*Gl*Rac signal observed by fluorescence microscopy, and the *Gl*Rac-positive structures were labeled with 0.2 µM DALGreen, as described above. Knockdown was confirmed by western blotting, as previously described (130). Halo-*Gl*Rac was detected using a mouse monoclonal anti-HaloTag antibody (IgG1; Promega, 1:1,000) and a goat anti-mouse IgG1 secondary antibody conjugated to Alexa Fluor 647 (Invitrogen, 1:2,500). Tubulin (loading control) was detected with a mouse monoclonal anti-acetylated tubulin antibody (clone 6-11B-1, IgG2b; MilliporeSigma, 1:3,000) and a goat anti-mouse IgG2b secondary antibody conjugated to Alexa Fluor 488 (Invitrogen, 1:2,500). Analyses were performed approximately 24 h after morpholino electroporation.

To assess the effect of constitutive activation of *Gl*Rac, *Gl*Rac^CA^ cells (tetracycline-inducible Q74L HA-*Gl*Rac) were grown in PE to log phase, then induced with 20 µg/mL tetracycline for 24 h. Halo-*Gl*Rac cells cultured under the same conditions served as controls. *Gl*Rac-positive compartments were labeled using GL50803_00137680-mNG, a cathepsin L-like protease that showed the most consistent expression across tested cathepsins.

### Live-cell imaging

Cells were chilled on ice for 20 min and vigorously shaken to detach from the culture tube. Cultures were then transferred to Attofluor cell chambers (Molecular Probes) and incubated at 37 °C for 1 h in a tri-gas incubator (Panasonic) set to 2% O_2_ and 5% CO_2_. To label Halo-tagged proteins, cells were mixed with 40 nM Janelia Fluor 646 HaloTag Ligand (Promega). Before imaging, chambers were washed three times with pre-warmed HBS.

For experiments involving morpholino knockdown or constitutively active *Gl*Rac, in which cells showed impaired attachment, treated cells were placed on ice for 20 min, centrifuged at 700 × g for 7 min, washed in cold HBS, and then transferred to coverslips pre-coated with 0.1% poly-L-lysine (MilliporeSigma). They were subsequently mounted on slides for imaging.

Imaging was performed using a DeltaVision Elite deconvolution microscope (GE) equipped with a 60×, 1.42 NA objective and a PCO Edge 5.4 sCMOS camera (PCO-TECH Inc.), using DIC, FITC, and Cy5 filter sets. Image deconvolution was carried out using SoftWoRx software (Applied Precision, API). For time-course experiments, imaging was conducted in a temperature and gas-controlled Bold Line stage-top incubator (Okolab) set to 37 °C, 2% O_2_, and 5% CO_2_.

### Time-lapse microscopy

To observe *Gl*Rac-positive compartment formation, Halo-*Gl*Rac cells were cultured for 48-72 h in PE and prepared in the Attofluor cell chambers as described above. To restrict motility, attached cells were overlaid with 1.5% ultra-low gelling temperature agarose (MilliporeSigma), prepared by dissolving 0.09 g of the polymer in 1.4 mL Milli-Q H_2_O, followed by incubation at 60 °C for 15 min (133). Next, 150 µL of 10× TYDK salts and 4.5 mL of sterile-filtered, nutrient-depleted medium obtained from 72-96 h PE cultures were added to the mixture. The final mix was kept at 37 °C until use. After aspirating the culture medium and washing the chambers with pre-warmed nutrient-depleted PE, 1 mL of the agarose mixture was gently added on top of the cells. Chambers were left at room temperature for 15 min in a GasPak EZ anaerobe pouch (BD Biosciences), then transferred to 4 °C for 5 min to allow the overlay to solidify. Imaging was performed under temperature and gas-controlled conditions as described in the live-cell imaging section.

To monitor compartment clearance, Halo-*Gl*Rac cells from 72 h PE cultures were attached to Attofluor chambers as described above. Following the attachment, cells were incubated with fresh TYDK for 10 min prior to agarose overlay. In this case, the agarose solution was prepared as above but mixed with 4.5 mL of fresh TYDK instead of nutrient-depleted medium. Imaging was performed under the same conditions as described for live-cell imaging.

### Transmission Electron Microscopy

Encysting trophozoites were processed using a protocol modified from previous work (134). Briefly, cells were allowed to attach to ACLAR sheets (Ted Pella) in 8-well plates for 1 h at 37 °C in a tri-gas incubator (2% O_2_, 5% CO_2_). Adherent cells were fixed with 2% paraformaldehyde (Ted Pella) and 2.5% glutaraldehyde (EMS) in 0.1 M sodium cacodylate buffer (EMS), pH 7.4, supplemented with 2 mM CaCl_2_ (MilliporeSigma). After rinsing with buffer, cells were treated sequentially with 2% osmium tetroxide (EMS) and 1.5% potassium ferrocyanide (MilliporeSigma), 1% thiocarbohydrazide (MilliporeSigma), a second 2% osmium tetroxide treatment (EMS), 2% uranyl acetate (EMS), and Walton’s lead aspartate (MilliporeSigma), with water rinses between each step. Samples were then dehydrated through ethanol (Decon) and acetone (EMS) and embedded in a hard formulation of EMBed 812 (EMS). Thin sections (60-80 nm) were collected on carbon-Formvar-coated grids and examined using a CM100 transmission electron microscope (Philips) equipped with a Morada camera (Olympus).

Nutrient-depleted trophozoites from 72 h cultures were processed similarly. After aldehyde fixation, cells were rinsed with buffer, quenched with 20 mM glycine (MilliporeSigma) in buffer, rinsed with buffer, and then treated with 2% osmium tetroxide (EMS) followed by 2% uranyl acetate (EMS), with water rinses between steps. Samples were dehydrated and embedded as described above. Grids were additionally contrasted with UranyLess (EMS) and lead citrate (EMS) before imaging.

### Image analysis

ImageJ version 1.54f (135) was used to process the images and measure the size of the *Gl*Rac-positive compartments. Figures were assembled in Adobe Illustrator CS6.

### Statistical analysis

All statistical analyses were performed using GraphPad Prism 10 for Windows (version 10.5.0). Paired two-tailed t-tests were used for comparisons in experiments where matched samples were analyzed. For comparisons of *Gl*Rac-positive structures per cell and compartment size across different conditions (displayed as violin plots), two-tailed Welch’s t-tests were used to account for unequal variances. P-values ≤ 0.05 were considered statistically significant. Significance levels are indicated as follows: *P ≤ 0.05, **P ≤ 0.01, ***P ≤ 0.001, ****P ≤ 0.0001. All microscopy images are representative of at least two independent experiments with comparable results.

## Data Availability

All study data are included in the article, Supporting information, or Dryad DOI: 10.5061/dryad.bzkh189sc

## Acknowledgements

The authors thank Dr. Justin Kollman for invaluable discussions and feedback on the manuscript and Dr. Vernon B. Carruthers for thoughtful discussions and for generously providing the BODIPY-LHVS cathepsin probe. We also thank the undergraduate students of the 2024 BIOL 402 course for designing and generating the *Gl*ATG constructs. This work was supported by National Institutes of Health Grants R01AI168417 and R21AI159035 (to A.R.P.).

## Supporting Information

**S1 Fig.**
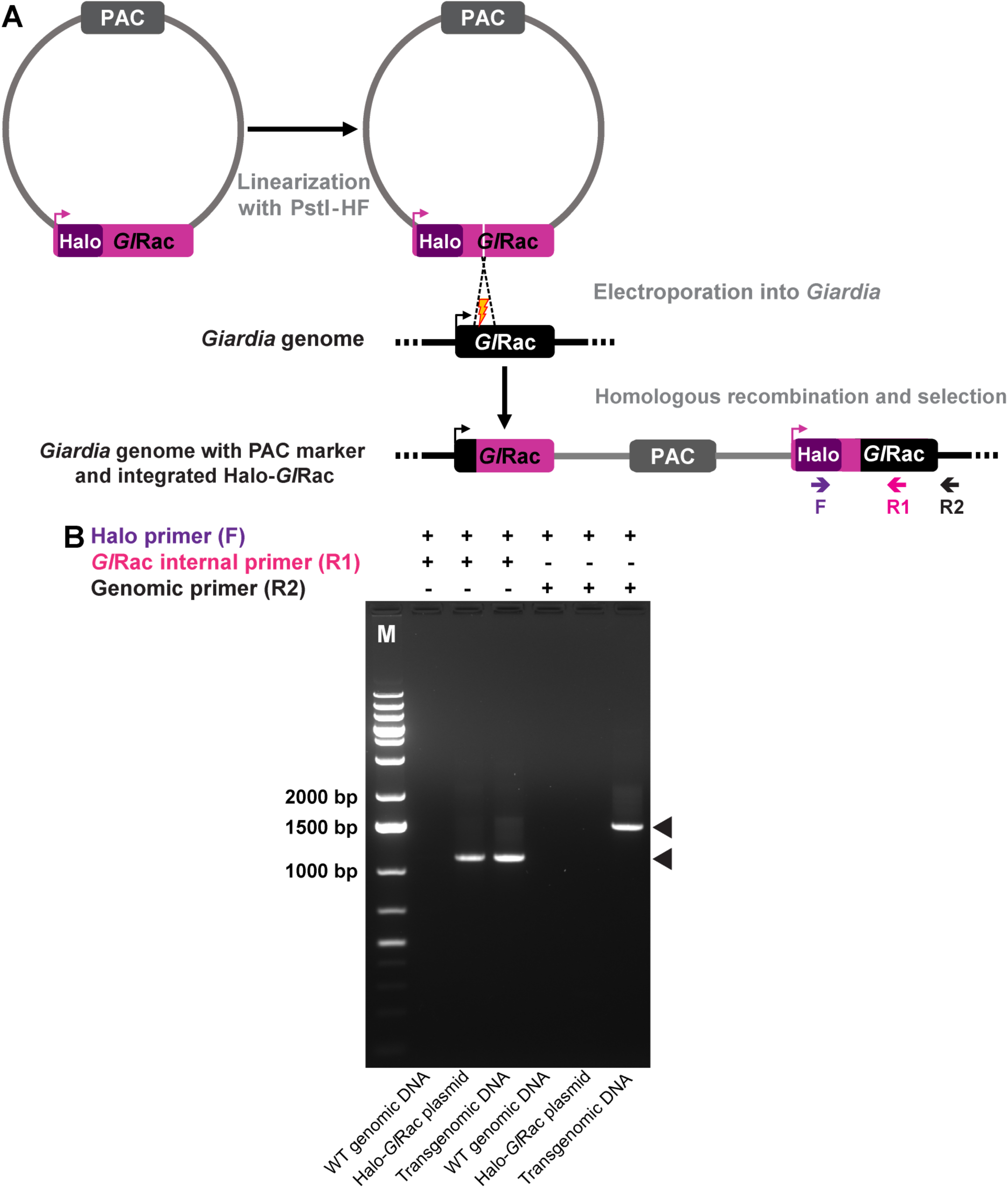
Strategy for Halo-*Gl*Rac genomic integration and validation by PCR. **(A)** Schematic representation of the plasmid components, integration strategy, and primer positions (F, R1, and R2) used to verify genomic integration of Halo-*Gl*Rac. The lightning icon represents a random double-strand DNA break in the *Gl*Rac locus. **(B)** Agarose gel showing PCR products confirming integration of the Halo-*Gl*Rac construct into the *Giardia* genome. PCR was performed using a common forward primer (Halo F) that anneals to the HaloTag sequence (absent from the *Giardia* genome) and two reverse primers: R1 (*Gl*Rac internal) and R2 (genomic). M: Thermo Scientific GeneRuler 1 kb Plus DNA Ladder. Black arrowheads indicate the expected PCR bands.

**S2 Fig.**
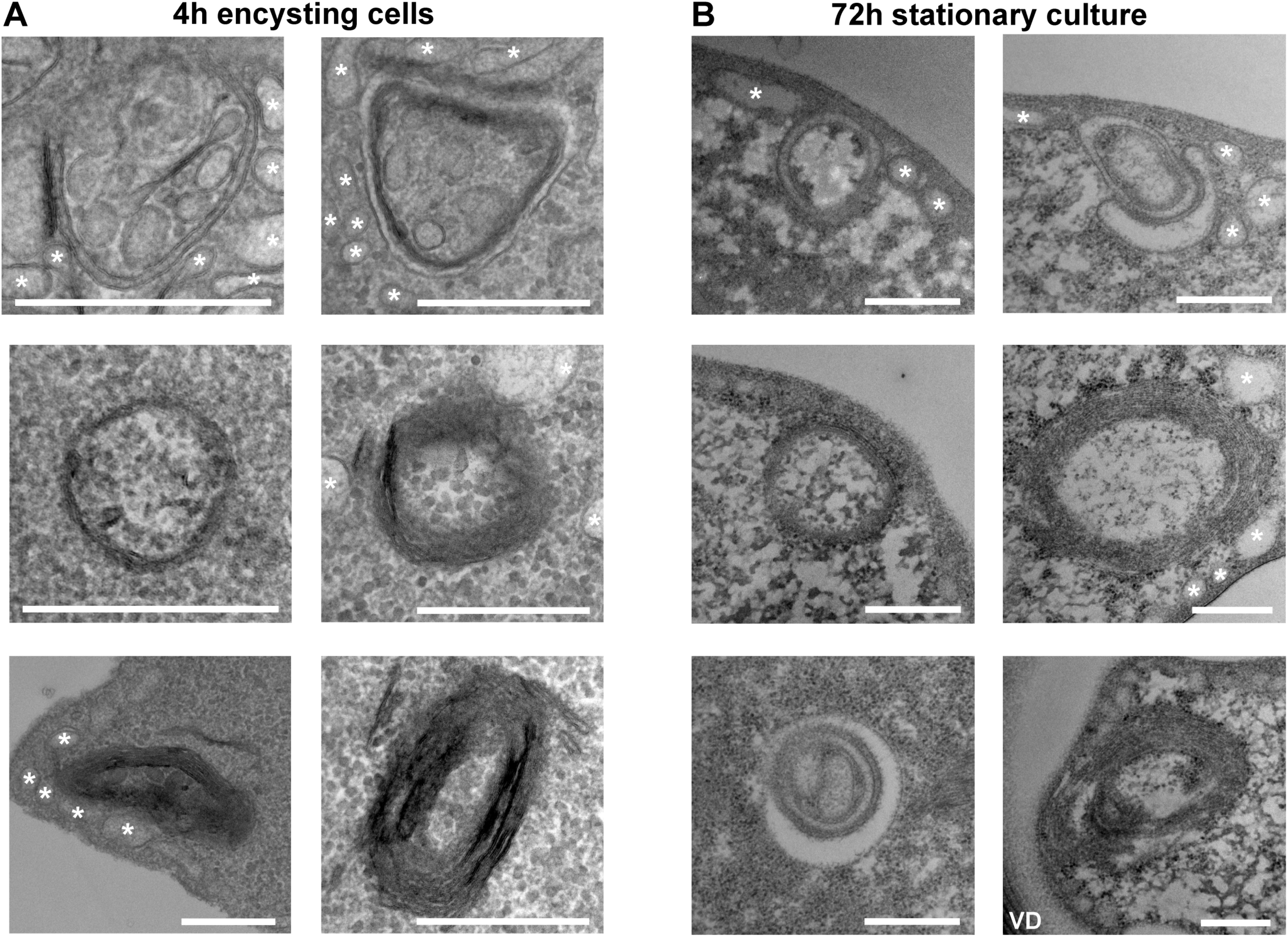
Double-membrane structures in starved cells display ultrastructural features similar to those observed during encystation. TEM images of 4 h encysting trophozoites **(A)** and trophozoites cultured for 72 h **(B)** reveal autophagosome-like structures with double-membrane morphology and multilamellar organization. Their size, morphology, and location are consistent with the *Gl*Rac-positive compartments identified by fluorescence microscopy, indicating that structures formed during nutrient depletion exhibit membrane architectures comparable to those observed during encystation. Asterisks indicate vesicles consistent with peripheral vacuoles (PVs). VD, ventral disc, which demarcates the bare area region where *Gl*Rac-positive compartments commonly form (see Fig 4A and S1 Movie). Scale bar, 0.5 µm.

**S3 Fig.**
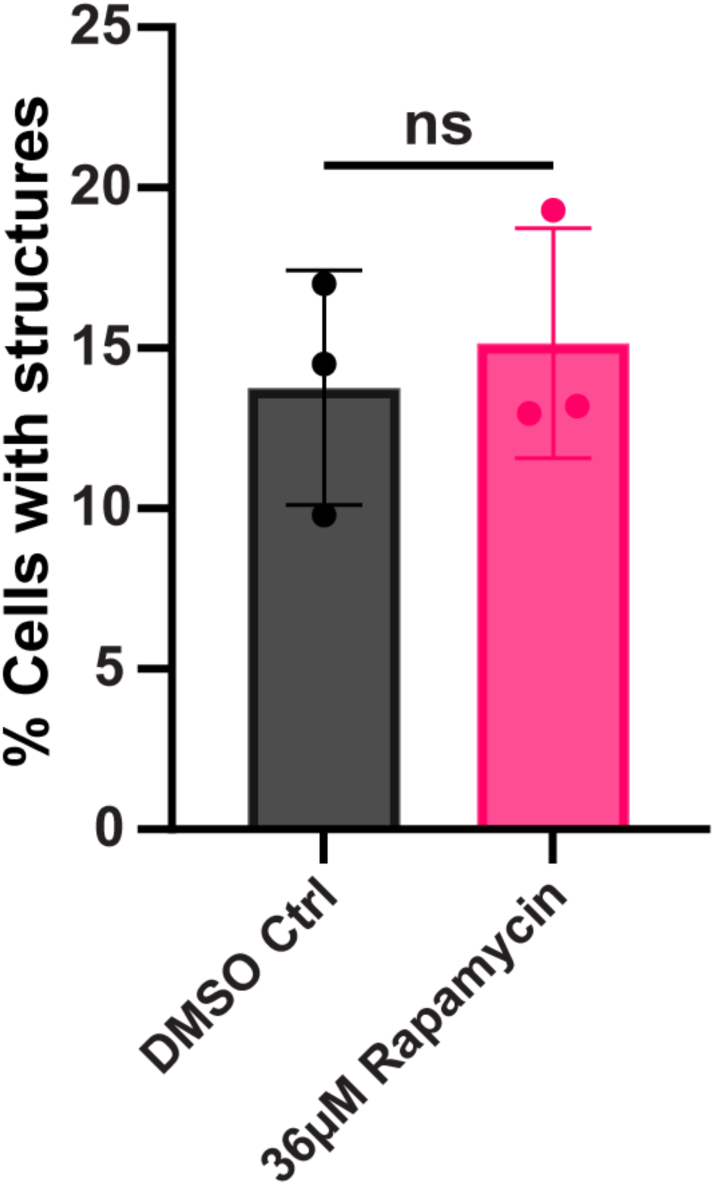
Rapamycin treatment does not significantly affect the abundance of *Gl*Rac-positive structures. Quantification of Halo-*Gl*Rac cells harboring *Gl*Rac-positive structures after 48 h of culture, followed by 2 h treatment with 36 µM rapamycin or DMSO (vehicle). For each condition, n > 380 cells. Data are presented as mean ± SD from three independent experiments. P-values were calculated using paired two-tailed t-tests (ns, not significant, P > 0.05).

**S4 Fig.**
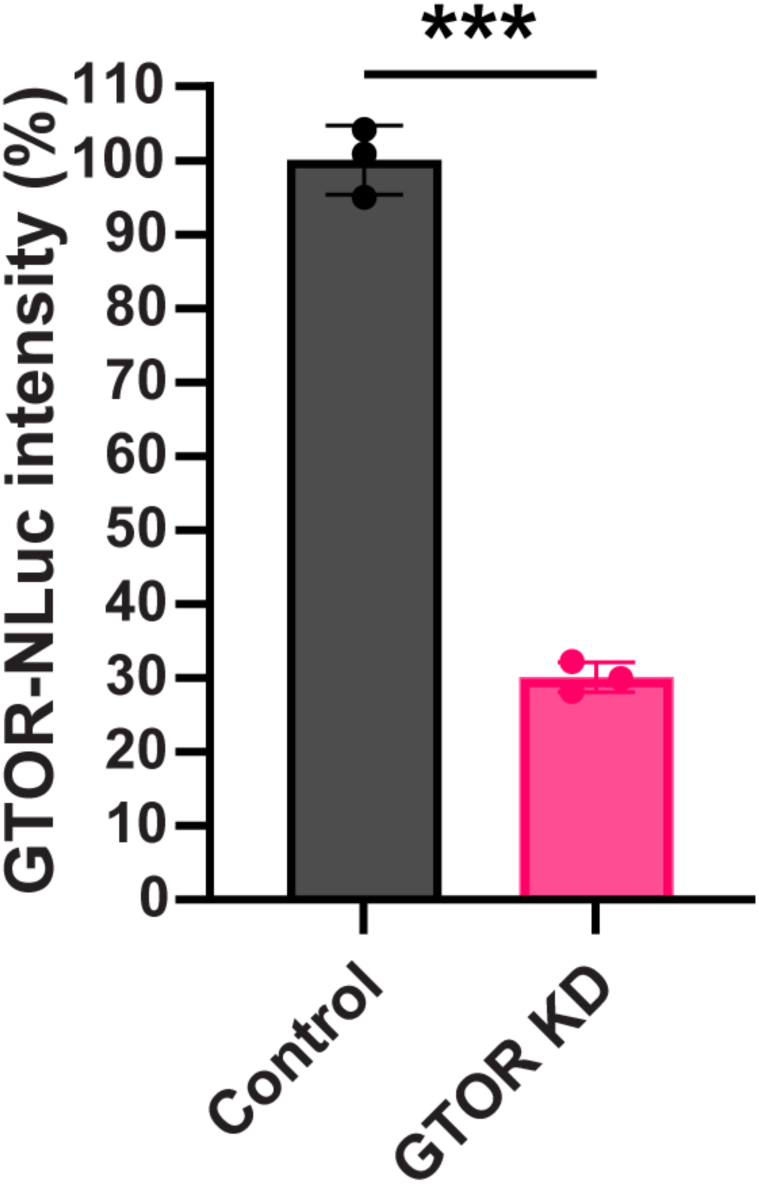
Translation-blocking anti-GTOR morpholino reduces GTOR levels by 69.9% relative to the standard control. Relative expression of endogenously tagged GTOR-NLuc after 24 h of transfection with either a translation-blocking morpholino targeting GTOR (GTOR KD) or a nonspecific control (control). Data are presented as mean ± SD from three experimental replicates. P-values were calculated using two-tailed t-tests (***P ≤ 0.001).

**S5 Fig.**
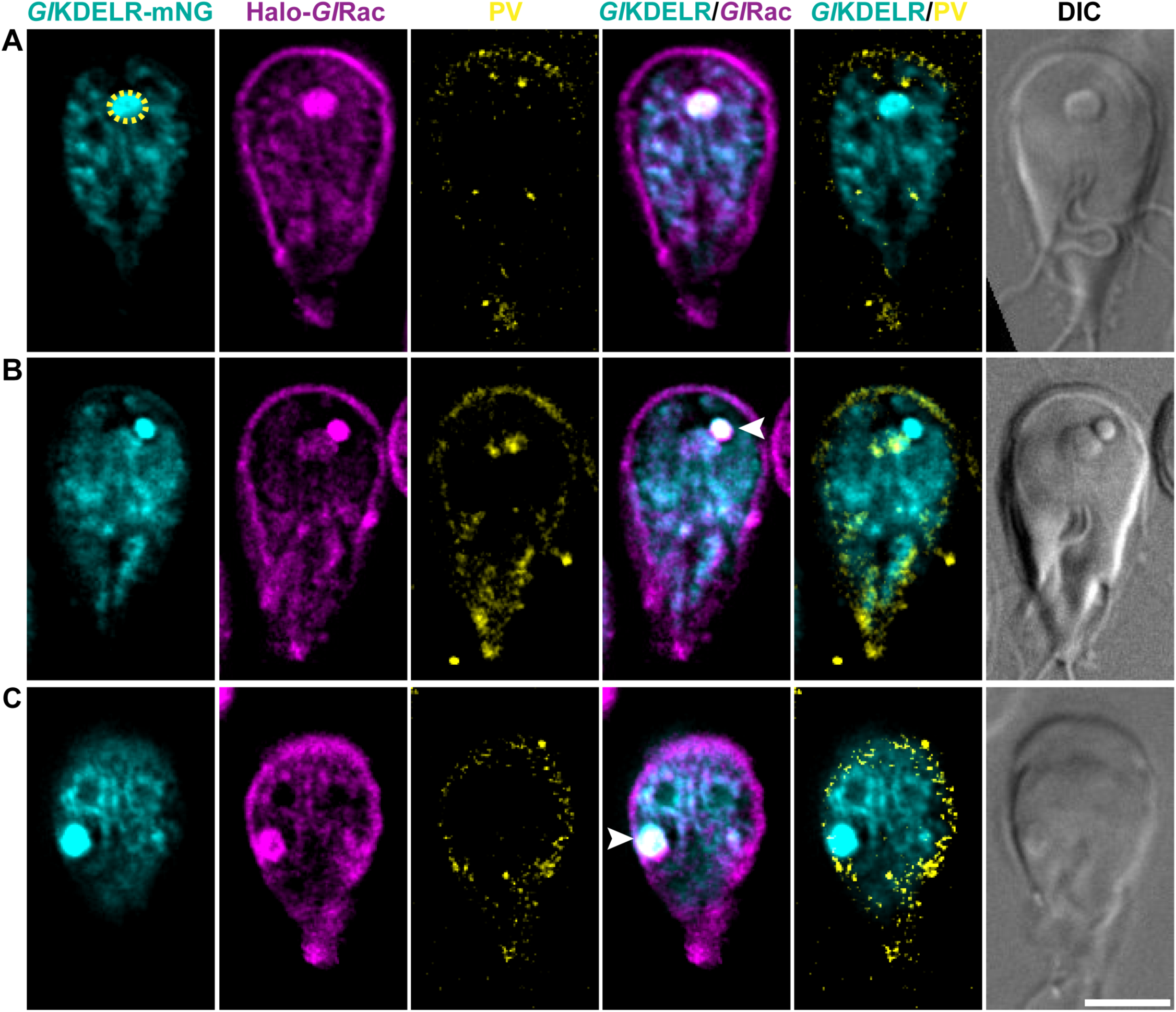
*Gl*KDELR is enriched in a ventral ER region and colocalizes with *Gl*Rac-positive organelles. Cellular distribution of *Gl*KDELR-mNG (cyan), Halo-*Gl*Rac (magenta), and Dextran Texas Red (yellow; PV marker). **(A)** *Gl*KDELR labels the endoplasmic reticulum throughout the cell but is enriched in a ventral region adjacent to the bare area (yellow dashed circle), where Halo-*Gl*Rac also accumulates. Based on the enrichment of the ER-associated protein *Gl*KDELR, this region is referred to as the ventral ER. Representative images showing colocalization of *Gl*KDELR-mNG and a *Gl*Rac-positive compartment (white arrowhead) in the ventral **(B)** and dorsal **(C)** portions of the cell. Scale bar, 5 µm.

**S6 Fig.**
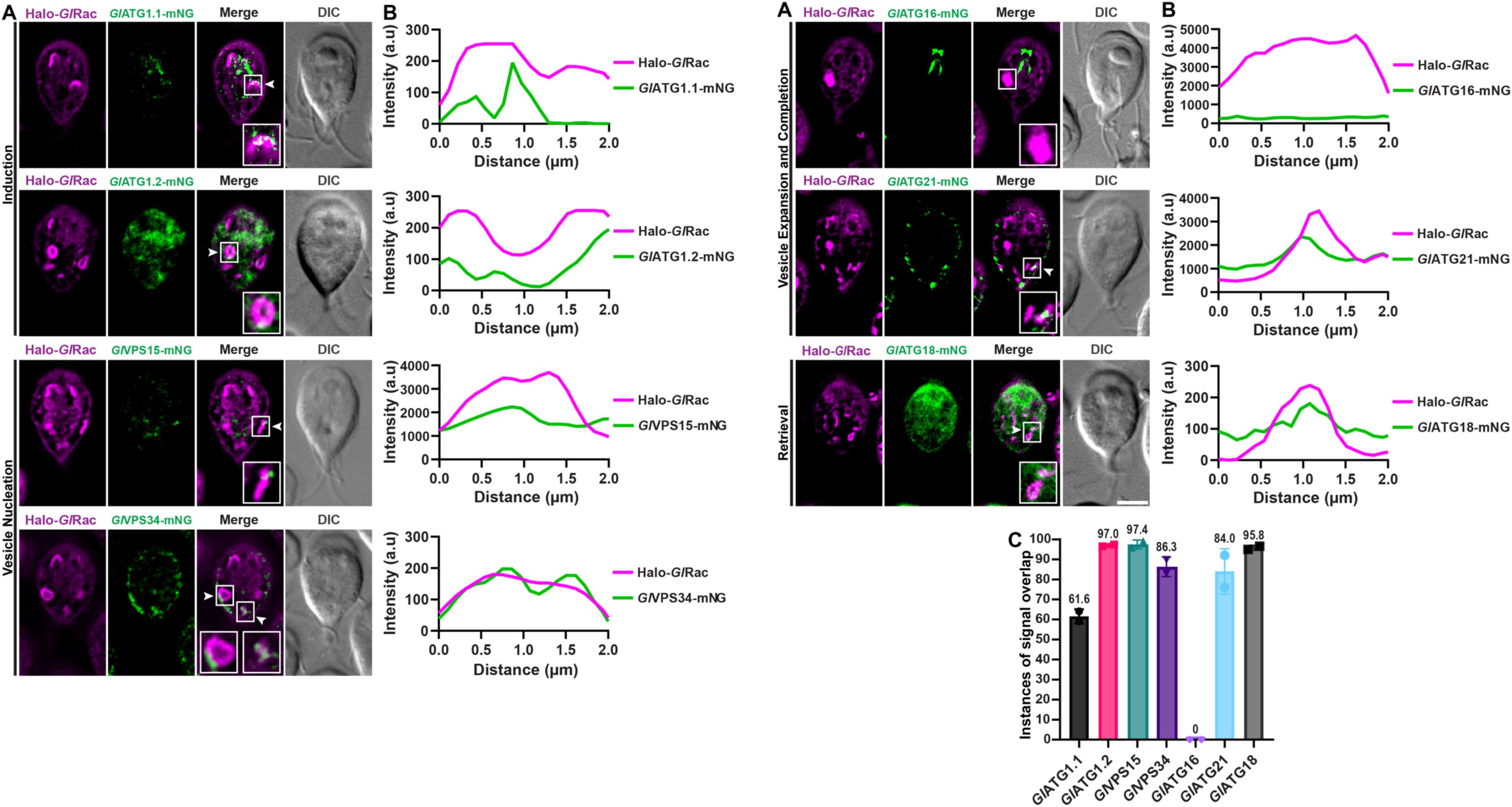
Cellular localization of *Giardia* ATG candidates identified by sequence similarity searches. **(A)** Halo-*Gl*Rac (magenta) cells expressing mNG-tagged putative *Gl*ATG1.1 (GL50803_0137719), *Gl*ATG1.2-like (GL50803_0017566), putative *Gl*VPS15 (GL50803_0113456), putative *Gl*VPS34 (GL50803_0017406), GL50803_0033762 (previously proposed as ATG16), putative *Gl*ATG21 (GL50803_0016957), or *Gl*ATG18-like (GL50803_0010822) were cultured for 72 h and imaged to assess localization. Representative images show the cellular distribution of each candidate protein (green). White arrowheads indicate examples of local signal overlap. Candidate proteins are grouped according to the stage of autophagy in which their proposed orthologs predominantly function (5). (**B)** Representative fluorescence intensity line scans across *Gl*Rac-positive structures illustrating local signal overlap between Halo-*Gl*Rac and the indicated candidate proteins. **(C)** Quantification of instances of signal overlap between Halo-*Gl*Rac-labeled structures and each candidate protein (>200 *Gl*Rac-positive structures analyzed from two independent experiments). Scale bar, 5 µm.

**S7 Fig.**
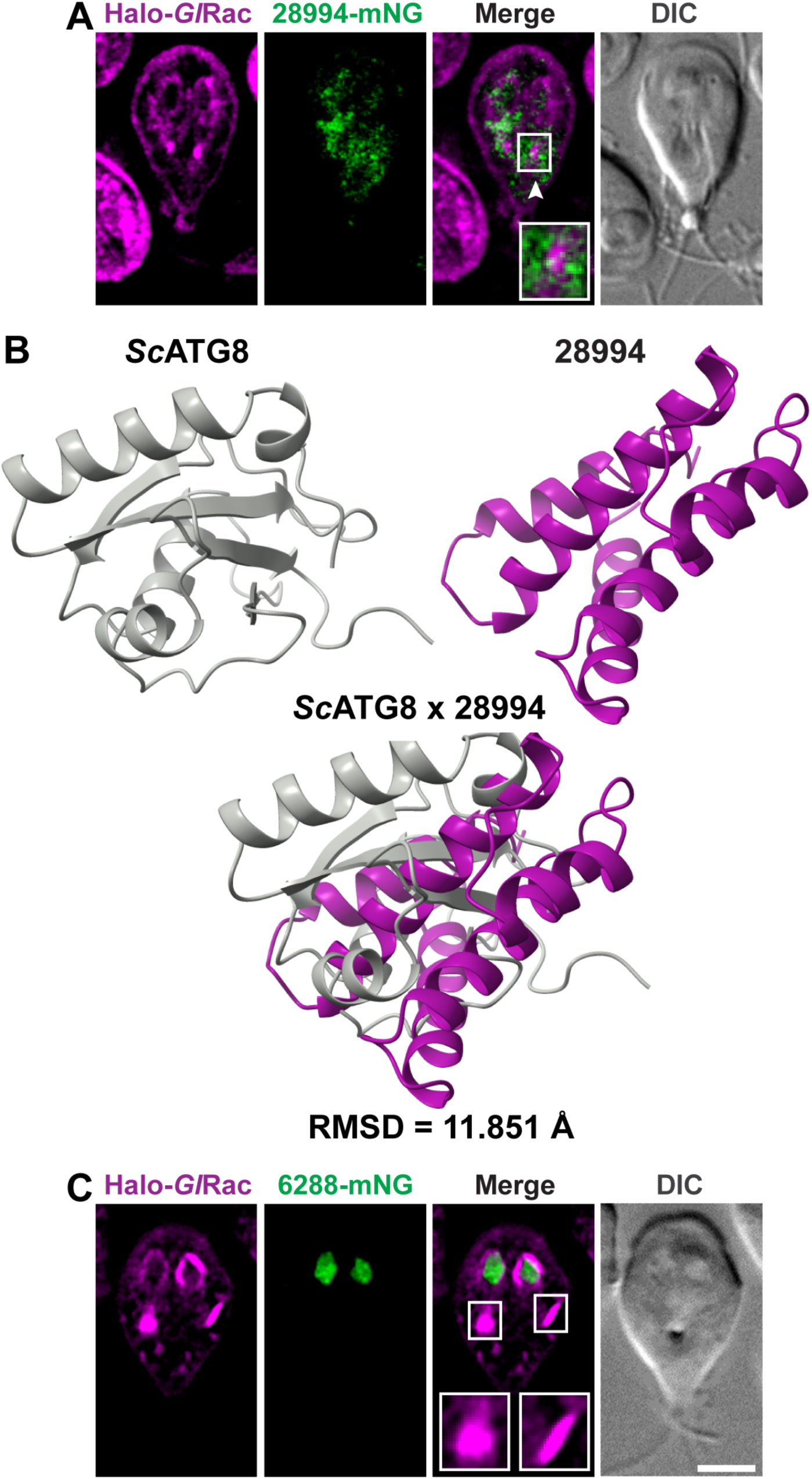
Evaluation of proposed *Giardia* ATG7 and ATG8 candidates lacking detectable sequence similarity. **(A)** Cellular distribution of GL50803_0028994-mNG (green; proposed ATG8 ortholog) in the Halo-*Gl*Rac (magenta) strain cultured for 72 h. The white arrowhead highlights a region of signal overlap. **(B)** Structural alignment of *S. cerevisiae* ATG8 (*Sc*ATG8, gray; UniProt ID: P38182) and GL50803_0028994 (magenta; UniProt ID: A8BQ35). Predicted structures were generated using AlphaFold and aligned in ChimeraX 1.8. The RMSD value is reported in Ångström (Å). **(C)** Cellular distribution of GL50803_006288-mNG (green; proposed ATG7 ortholog) in the Halo-*Gl*Rac (magenta) strain cultured for 72 h. No overlap was observed between Halo-*Gl*Rac-labeled structures and 6288-mNG (n > 200 cells from two independent experiments). Scale bar, 5 µm.

**S8 Fig.**
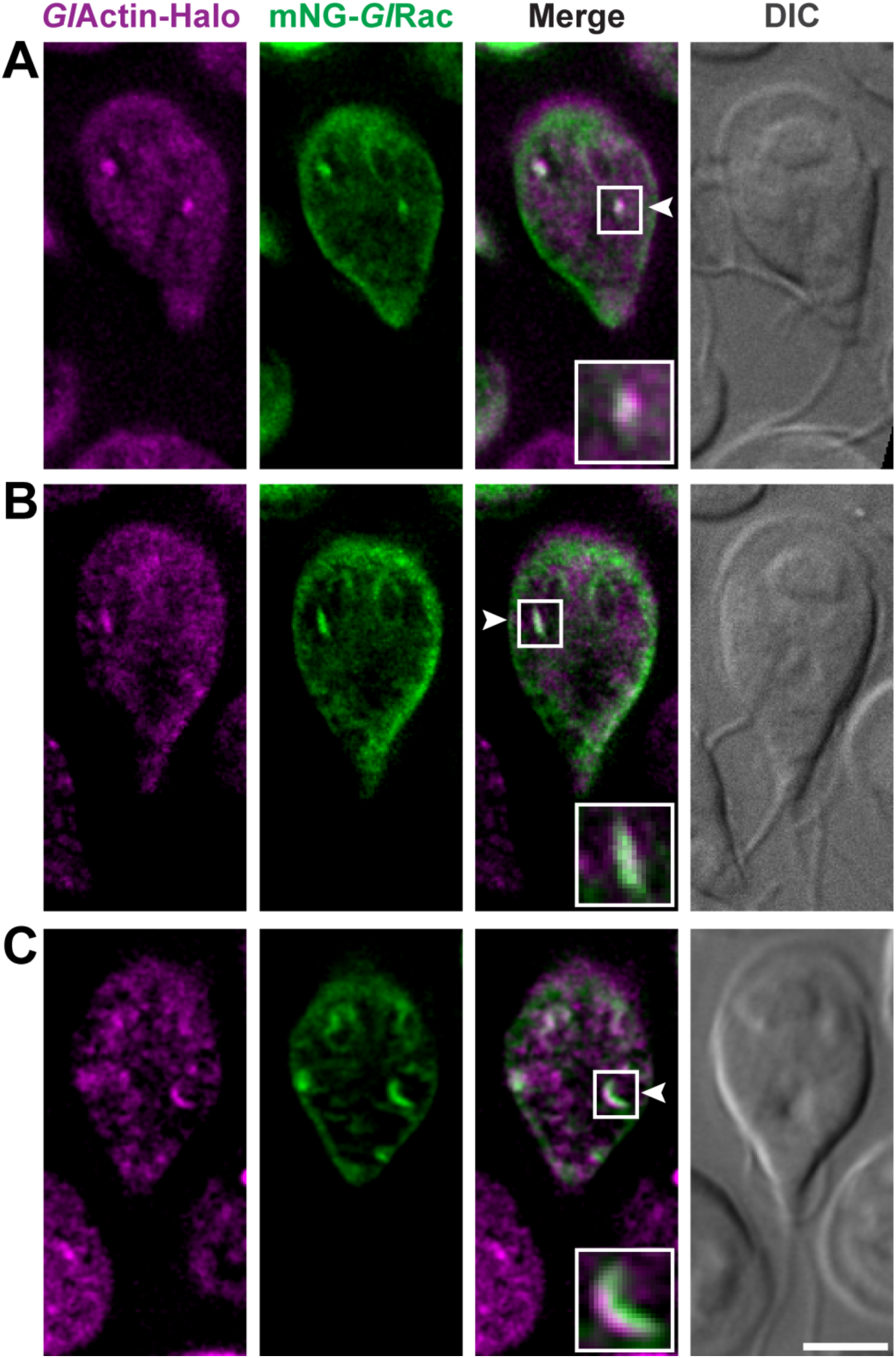
Actin associates with *Gl*Rac-positive intermediate structures. Representative live-cell images of trophozoites expressing *Gl*Actin-Halo and mNG-*Gl*Rac. Actin-Halo localizes to *Gl*Rac-positive structures at multiple stages of compartment formation, including short linear intermediates **(A)**, extended linear intermediates **(B)**, and cup-shaped intermediates **(C)**. Insets show enlarged views of the indicated regions. White arrowheads indicate close association of actin with *Gl*Rac-positive structures during the early stages of compartment formation. Scale bar, 5 µm.

**S9 Fig.**
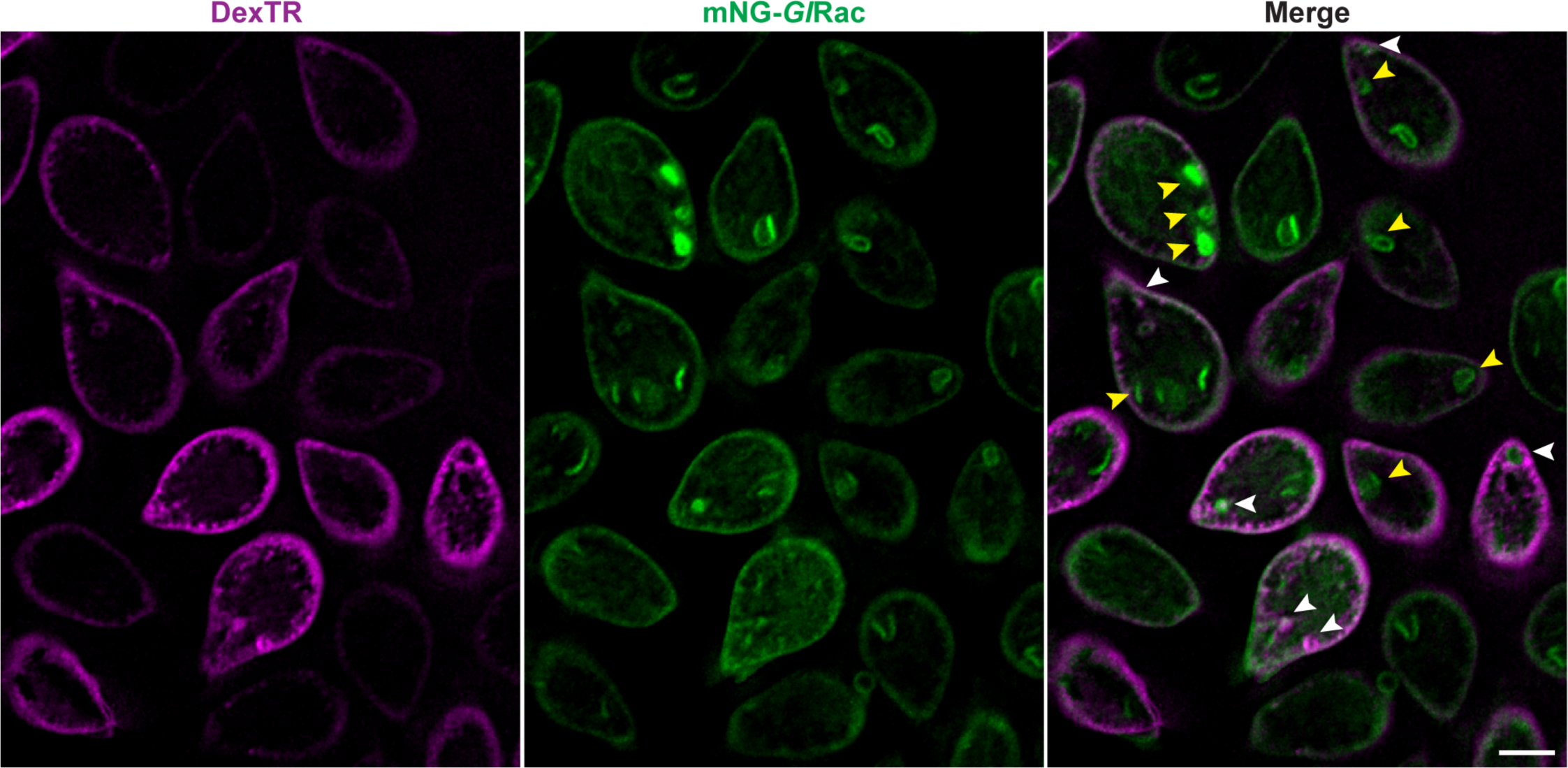
*Gl*Rac-positive organelles display heterogeneous association with Dextran Texas Red-labeled peripheral vacuoles. mNG-*Gl*Rac cells cultured for 48 h and then treated with 1 mg/mL Dextran Texas Red (DexTR; magenta) for 24 h. Representative images showing *Gl*Rac-positive structures associated with DexTR-labeled peripheral vacuoles (PVs), including structures surrounded by or containing DexTR-positive material (white arrowheads), as well as structures with little or no detectable DexTR association (yellow arrowheads). Scale bar, 5 µm.

**S10 Fig.**
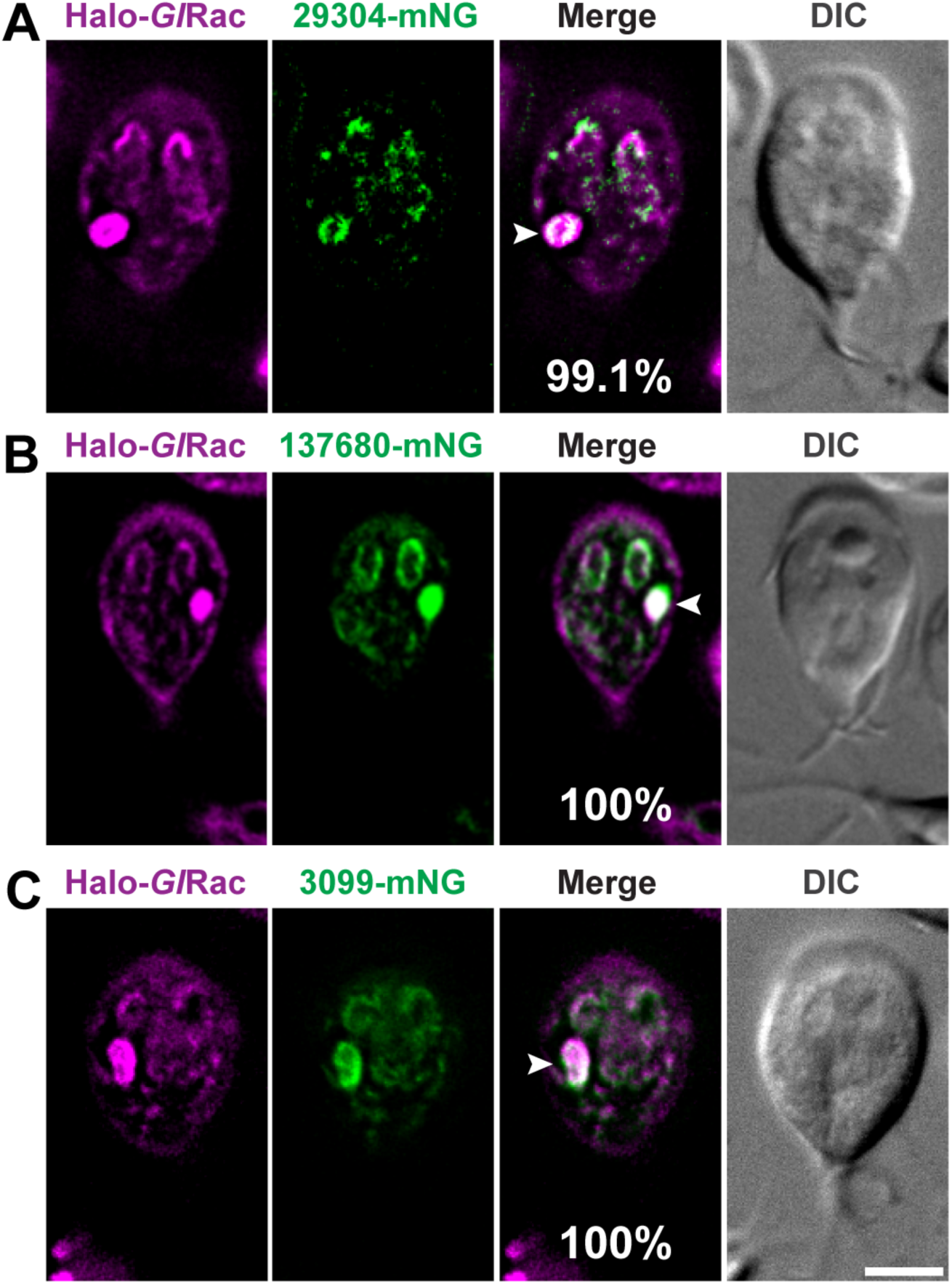
Additional cathepsins recruited to *Gl*Rac-positive organelles. **(A-C)** Cellular distribution of cathepsin B (GL50803_0029304-mNG; green) **(A)** and two cathepsin L-like proteases, GL50803_00137680-mNG **(B)** and GL50803_003099-mNG **(C)**, in the Halo-*Gl*Rac (magenta) strain cultured for 72 h. Among these, GL50803_00137680-mNG showed the most consistent expression and was used to label these compartments in experiments where Halo-*Gl*Rac could not be visualized (Fig 7E). White arrowheads indicate colocalization. The percentages of cells showing colocalization between cathepsins and *Gl*Rac-positive structures are indicated in the figure. For each condition, n > 200 cells from two independent experiments. Scale bar, 5 µm.

**S11 Fig.**
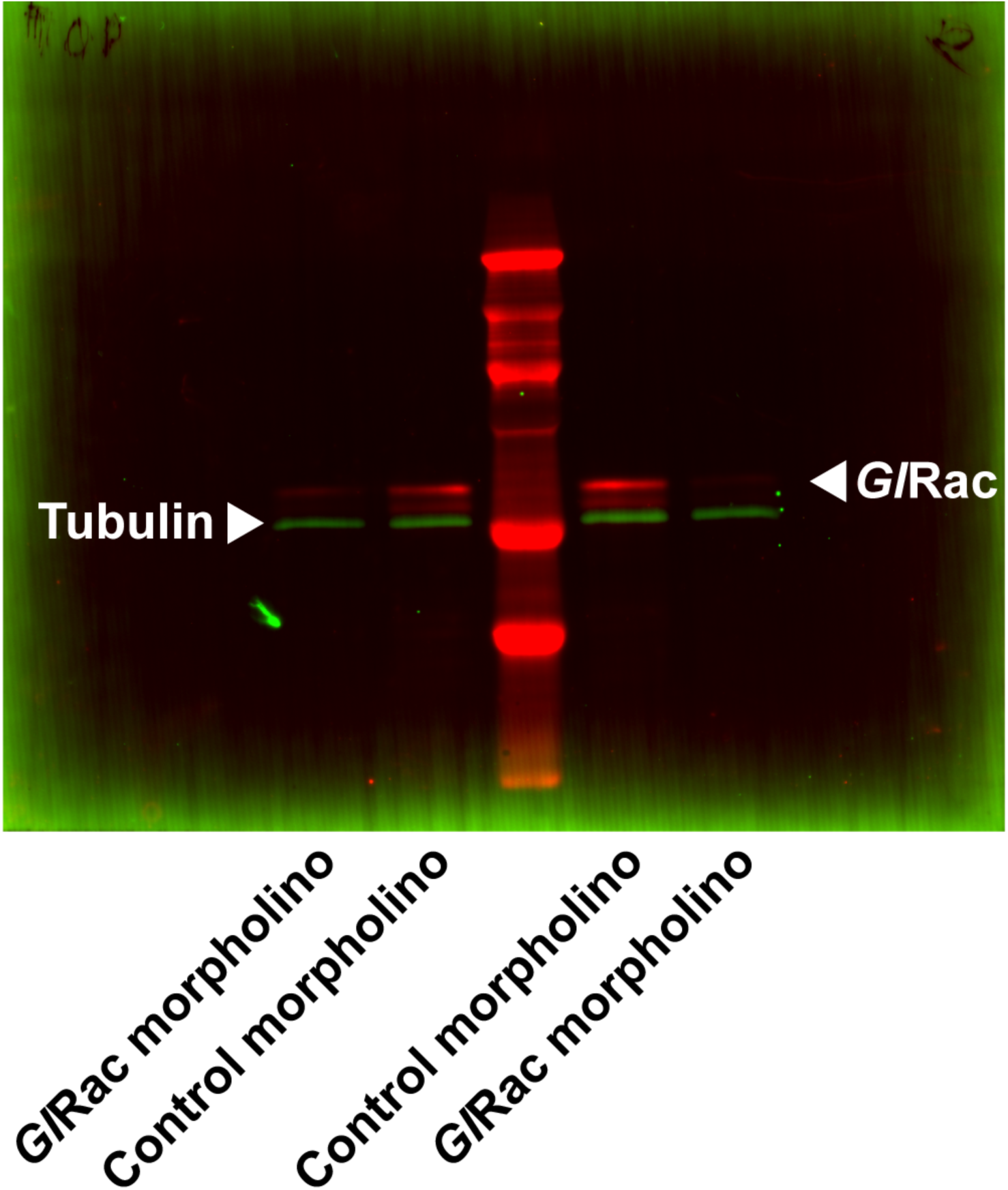
Translation-blocking anti-*Gl*Rac morpholino reduces *Gl*Rac levels by 70.3% relative to the standard control. Immunoblots were performed 24 h after morpholino treatment, matching the timing of live-cell experiments. Membranes were immunoblotted with anti-Halo and anti-tubulin antibodies, and *Gl*Rac levels (anti-Halo) from endogenously tagged cell lines were normalized to tubulin as a loading control. This depletion confirms the efficacy of this previously validated morpholino (39,88).

**S12 Fig.**
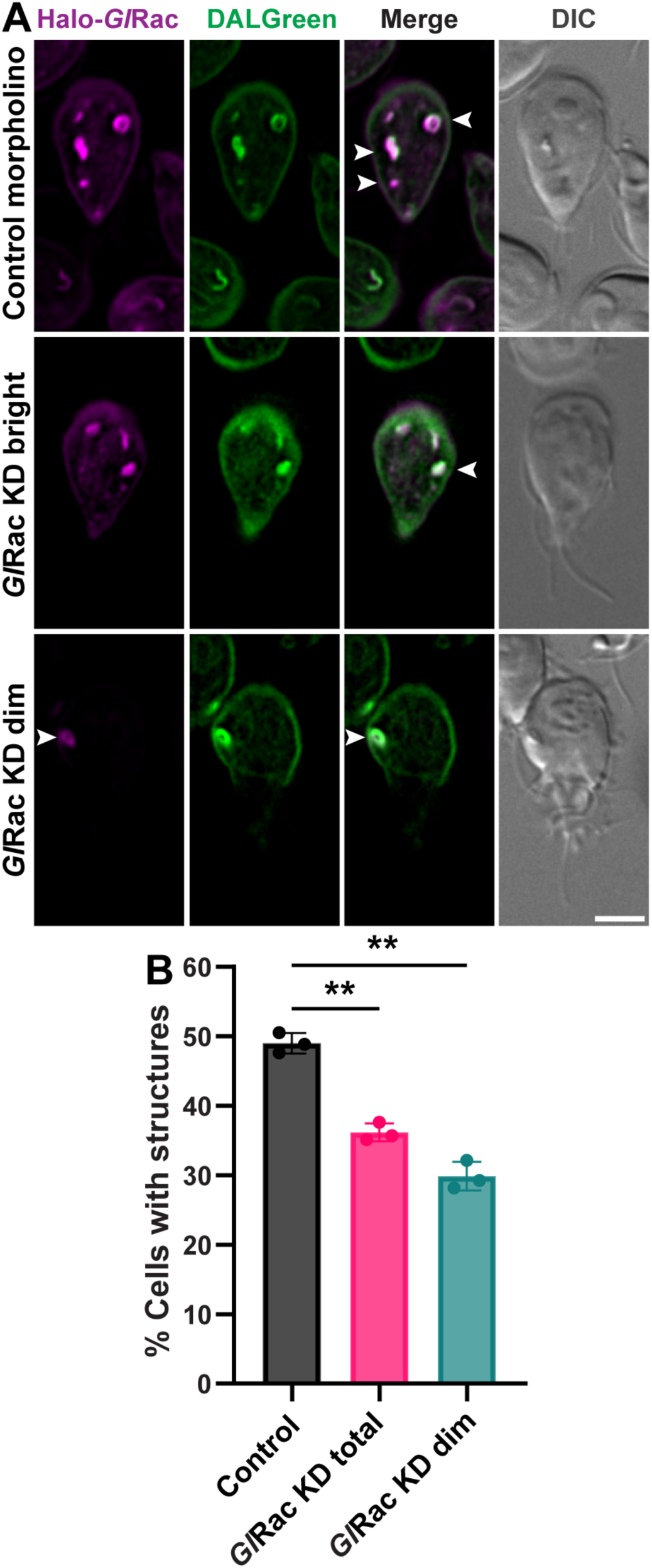
Variable morpholino penetrance observed after *Gl*Rac knockdown. **(A)** Representative images of Halo-*Gl*Rac^MS^ cells 24 h after transfection with either a nonspecific control morpholino (Ctrl morpholino) or a translation-blocking morpholino targeting *Gl*Rac (*Gl*Rac KD). The *Gl*Rac KD population includes cells with a high Halo-*Gl*Rac signal and typically normal morphology (*Gl*Rac KD bright) and cells with reduced signal and morphological defects (*Gl*Rac KD dim). White arrowheads indicate colocalization of Halo-*Gl*Rac-labeled organelles with DALGreen. Scale bar, 5 µm. **(B)** Expanded quantification of Halo-*Gl*Rac^MS^ cells harboring *Gl*Rac-positive structures, including *Gl*Rac KD total (bright + dim) cells. This complements Fig 7B, which only includes the control morpholino and *Gl*Rac KD dim subsets. For each condition, n > 260 cells. Data are presented as mean ± SD from three independent experiments. P-values were calculated using paired two-tailed *t*-tests (**P ≤ 0.01).

**S1 Dataset. Sequence-based identification of *Giardia lamblia* ATG candidates and cathepsins analyzed in this study.** Comprehensive list of putative *Giardia lamblia* ATG genes identified through prior bioinformatic analyses or reciprocal BLAST searches, along with cathepsins analyzed in this work. All candidates were C-terminally tagged and expressed in the Halo-*Gl*Rac background for localization studies. ATG candidates and cathepsins are provided in separate spreadsheet tabs.

**S2 Dataset. Primers and constructs used in this study.** List of primers used to PCR-amplify *Giardia lamblia* coding sequences, together with associated construct details.

**S1 Movie. Formation of a *Gl*Rac-positive compartment at the bare area.** Live-cell imaging of a trophozoite expressing Halo-*Gl*Rac (magenta) showing formation of a *Gl*Rac-positive compartment adjacent to the bare area of the ventral disc. The compartment initially appears as a faint linear signal, progressively elongates and enlarges, and then transitions into curved and circular morphologies. Images were acquired every 5 min.

**S2 Movie. Formation of a *Gl*Rac-positive compartment in the dorsal area.** Live-cell imaging of a trophozoite expressing Halo-*Gl*Rac (red) showing formation of a *Gl*Rac-positive compartment in the dorsal region of the cell. The compartment initially appears as a faint linear signal, progressively elongates and enlarges, and then transitions into curved and circular morphologies. Images were acquired every 3 min.

**S3 Movie. Clearance of a *Gl*Rac-positive compartment.** Live-cell imaging of a trophozoite expressing Halo-*Gl*Rac (magenta) showing clearance of a *Gl*Rac-positive compartment. The compartment initially appears circular, then transitions to a curved morphology, progressively shrinks, and is ultimately reabsorbed by the cell. Images were acquired every 2 min.

